# Quantification of biases in predictions of protein-protein binding affinity changes upon mutations

**DOI:** 10.1101/2023.08.04.551687

**Authors:** Matsvei Tsishyn, Fabrizio Pucci, Marianne Rooman

## Abstract

Understanding the impact of mutations on protein-protein binding affinity is a key objective for a wide range of biotechnological applications and for shedding light on disease-causing mutations, which are often located at protein-protein interfaces. Over the past decade, many computational methods using physics-based and/or machine learning approaches have been developed to predict how protein binding affinity changes upon mutations. They all claim to achieve astonishing accuracy on both training and test sets, with performances on standard benchmarks such as SKEMPI 2.0 that seem overly optimistic. Here we benchmarked eight well-known and well-used predictors and identified their biases and dataset dependencies, using not only SKEMPI 2.0 as a test set but also deep mutagenesis data on the SARS-CoV-2 spike protein in complex with the human angiotensin-converting enzyme 2. We showed that, even though most of the tested methods reach a significant degree of robustness and accuracy, they suffer from limited generalizability properties and struggle to predict unseen mutations. Interestingly, the generalizability problems are more severe for pure machine learning approaches while physics-based methods are less affected by this issue. Moreover, undesirable prediction biases towards specific mutation properties, the most marked being towards destabilizing mutations, are also observed and should be carefully considered by method developers. We conclude from our analyses that there is room for improvement in the prediction models and suggest ways to check, assess and improve their generalizability and robustness.

## 1 Introduction

Proteins interact with each other to form complexes that perform a wide range of biological functions in the intra- and extracellular media, and are involved in key processes such as signal transduction, cell growth and proliferation, and cell apoptosis. It is therefore of fundamental interest to understand how amino acid substitutions impact on the ability of proteins to bind to their interacting partners. Such insights would shed light on pathogenic mechanisms since aberrant protein-protein interactions caused by deleterious variants are often central to Mendelian disorders and complex diseases such as cancer [1, 2, 3, 4]. From a biotechnological perspective, it would improve the design of drugs that modulate protein-protein interactions (PPIs), as targeting these is an established strategy in the treatment of disease [5, 6].

There are several experimental methods for estimating the impact of mutations on PPIs. Biophysical methods such as isothermal titration calorimetry allow in-depth estimation of protein binding thermodynamics [7]; in contrast, high-throughput screening assays such as yeast-two-hybrid systems only allow identification of binary PPIs but have the advantage of being applicable at a large scale [8]. However, given that all experimental approaches remain challenging, costly and time-intensive, there is room for computational methods which provide effective alternatives to predict and achieve better understanding of PPIs.

Over the last decade, many studies have been dedicated to the development of bioinformatics tools to predict the impact of mutations on protein-protein binding affinity (Δ*G*_*b*_), which is the thermodynamic descriptor of protein-protein interactions [9, 10, 11, 12, 13, 14, 15, 16, 17, 18, 19, 20, 21]. These tools are mainly based on structural features derived from experimentally characterized protein complexes and/or evolutionary data. These features are usually combined using standard machine learning techniques, but deep learning algorithms are starting to be used in predictor construction [20].

The first attempts to predict protein-protein binding affinity changes upon mutations (ΔΔ*G*_*b*_) were based on physical energy functions [22], with predictors such as Rosetta [9] (2002), FOLDEF [10] (2002) and DComplex [11] (2004). The lack of sufficiently large and standardized datasets of experimental ΔΔ*G*_*b*_ values prevented them from being trained directly on such data. For this reason, some of them (e.g. DComplex) were completely unsupervised, while others (e.g. Rosetta and FOLDEF) were trained on experimental values of protein stability changes upon mutations (ΔΔ*G*) reported in the ProTherm [23] dataset, with the assumption that physical properties of intraprotein interactions are transposable to interprotein interactions at the interface. In this case, experimental data were used only to parameterize the energy functions and to weight their individual contributions.

Now, the SKEMPI dataset [24, 25] fills this gap. It is considered as the gold standard for training and testing ΔΔ*G*_*b*_ predictors. Its first release in 2012, SKEMPI 1.0 [24], collected, curated, selected and standardized entries from literature searches and from already existing datasets (ASEdb [26], PINT [27], and [28]). This first release allowed the development of a generation of ΔΔ*G*_*b*_ predictors such as BeAtMuSiC [12] (2013), mCSM [13] (2014), MutaBind [14] (2016) and BindProfX [15] (2017). The large amount of collected experimental values enabled a more extensive use of machine learning methods (e.g. in mCSM), as well as leveraging other non-physical information to predict energy values. For instance, evolutionary information was extracted from homologous structures (in BindProfX) and sequences (in MutaBind).

The second SKEMPI release in 2019, SKEMPI 2.0 [25], increased the number of entries by more than a factor of two by adding new entries from literature and some more recent datasets (AB-Bind [29], PROXiMATE [30], dbMPIKT [31], and [32]). Moreover, it provided a more diverse set of mutations on a more diverse set of protein complexes. SKEMPI 2.0 allowed an even more extensive use of machine learning techniques and the development of a wider range of features, leading to a new generation of predictors, such as mCSM-PPI2 [16] (2019), MutaBind2 [17] (2020), SSIPe [18] (2020), SAAMBE-3D [19] (2020), NetTree [20] (2020) and mmCSM-PPI [21] (2021).

While these tools achieve good prediction accuracy on their respective training sets, the extent to which these results are generalizable to unseen data is one of the open issues in the field. Indeed, like all supervised machine learning methods, they are likely to suffer from undesirable biases towards the learning set, which often hinder the generalization of their predictions. One example of this problem is the bias towards destabilizing values of the folding free energy change upon mutations (ΔΔ*G*), which has been thoroughly analyzed in a series of investigations [33, 34, 35]. In summary, it has been shown that training protein stability predictors on the common experimental datasets that are dominated by destabilizing mutations leads to much better performance on destabilizing than on stabilizing mutations.

Although prediction biases have been studied for predictors of stability changes caused by mutations, they have not been for protein-protein affinity changes; yet having accurate and unbiased prediction tools of ΔΔ*G*_*b*_ values is crucial for a wide range of biotechnological applications. In this paper, we have systematically quantified possible biases in state-of-the-art protein-protein ΔΔ*G*_*b*_ prediction methods. More precisely, we evaluated their predictions on a set of mutations with experimentally measured ΔΔ*G*_*b*_ values taken from [25], and on high-throughput data on the binding between the human angiotensin-converting enzyme 2 (ACE2) and the receptor binding domain (RBD) of the SARS-CoV-2 spike protein taken from [36]. After an analysis of the methods’ performances, we suggest strategies to limit and correct possible biases and thus to further improve the methods’ generalizability and scores.

## 2 Methods

### 2.1 Protein-protein binding affinity change upon mutations

The thermodynamic protein-protein binding affinity Δ*G*_*b*_ is a measure of the strength of a PPI and is defined using the Gibbs free energy:

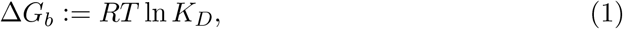

where *R* is the Boltzmann constant, *T* the absolute temperature (in K) and *K*_*D*_ the equilibrium dissociation constant of the PPI. We use the convention that the stronger the interaction, the more negative the value of Δ*G*_*b*_, and express it in kcal/mol.

Under the action of a mutation, we define the binding affinity change as:

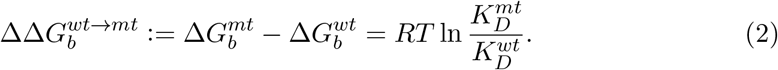

where *wt* refers to the wild-type complex and *mt* to the mutant. Thus, positive ΔΔ*G*_*b*_ values correspond to mutations that destabilize the complex and negative values to stabilizing mutations. Since binding affinity is a thermodynamic state function, mutating from a wild-type complex to a mutant complex and then mutating back results in no change in ΔΔ*G*_*b*_, which is expressed by the following equation:

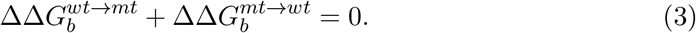

We will refer to this property as the symmetry property.

In what follows, we will call “direct mutation” a mutation that goes from the wild-type to the mutant complex. Conversely, we will call “reverse mutation” a mutation that goes from the mutant to the wild-type complex. Note that the terms wild-type, mutant, direct and reverse are defined with respect to the proteins that are part of our datasets and do not necessarily have a biological interpretation.

### 2.2 Defining protein-protein interfaces

The relative solvent accessibility (RSA) of a residue in a three-dimensional (3D) structure is defined as the ratio (in %) of its solvent-accessible surface area in the structure and in an extended tripeptide Gly-X-Gly [37]. We calculated them using our in-house software MuSiC [38] (which uses an extension of the DSSP algorithm [39]), available on the dezyme.com website. We distinguished between interactant-RSA (iRSA) and complex-RSA (cRSA), which correspond to the RSA calculated from the structure containing solely the considered interactant and from the structure containing the complex with both interactants, respectively. We defined the RSA change upon binding as ΔRSA := iRSA − cRSA; it measures how much the protein-protein interaction changes the solvent accessibility of a residue. A residue is considered to be in the protein-protein interface if its ΔRSA is greater than 5%.

### 2.3 Datasets of binding affinity changes upon mutations

We considered two datasets. The first is based on the SKEMPI sets [24, 25], containing mutations in different protein-protein complexes of known 3D structure available in the Protein Data Bank (PDB) [40], whose ΔΔ*G*_*b*_ values have been measured experimentally using biophysical methods, performed by various laboratories. The number of characterized mutations in each protein typically ranges from a few to a few dozen, and reaches in rare cases a few hundred [41, 42]. These datasets yield relatively accurate ΔΔ*G*_*b*_ values but have the disadvantage of being unsystematic and of reflecting the specific interests of the authors in the choice of proteins and mutations.

The SKEMPI 2.0 dataset [25], contains 7085 entries and is the most comprehensive, well curated and diverse dataset of its kind. First, we discarded entries without ΔΔ*G*_*b*_ value and entries describing multiple mutations. We then aggregated all redundant entries (with the same mutation in the same PDB structure) by taking their average ΔΔ*G*_*b*_ value. To withdraw the dependency on the quality of the structures, we also dropped all mutations in low-resolution X-ray structures (resolution *>* 2.5°A) and in structures obtained by nuclear magnetic resonance spectroscopy. This defines our first benchmark dataset called S2536 which contains 2536 mutations in 205 different PDB structures.

The second dataset we considered contains affinity values obtained through deep mutagenesis experiments that systematically characterized all possible mutations in the receptor binding domain (RBD) of the SARS-CoV-2 spike glycoprotein in interaction with the human ACE2 receptor [36]. This dataset has the advantage of being systematic and therefore less biased. However, the measured values are not exact ΔΔ*G*_*b*_ but close correlates. From this set, we first discarded the mutations of the few residues located in the N- and C-terminal tails of the spike protein, as they are absent from the reference PDB structure 6M0J. We then identified the ACE2-RBD interface residues, of which there are 20, using the above RSA criterion. We focused on all 380 possible mutations of these 20 residues, to define our second benchmark dataset C380.

For both the S2536 and C380 datasets, considered by definition as direct mutations, we constructed the datasets of reverse mutations using the symmetry property Eq. (3) to assign a ΔΔ*G*_*b*_ value to each reverse mutation. When the distinction is required, we append the suffix -D to the name for a dataset of direct mutations, the suffix -R for a dataset of reverse mutations and the suffix -DR for a dataset of both direct and reverse mutations (e.g., S2536-D, S2536-R and S2536-DR).

The datasets S2536 and C380 are available at https://github.com/3BioCompBio/DDGb_bias.

### 2.4 Protein 3D structures

For predicting direct mutations in the S2536 set, we used the PDB structures of the protein complexes that have been collected in the SKEMPI 2.0 database, as they were curated to be as close as possible to the protein complexes on which the measurements were made. For direct mutations in the C380 set, we used the experimental 3D structure of the ACE2-RBD complex with PDB ID 6M0J [43], as referenced in [36].

For reverse mutations, we modelled the mutant complexes using the comparative modeling software MODELLER [44] with default parameters and the wild-type structures as templates. MODELLER reconstructs the side chain of the mutated residue, then slightly rearranges the backbone and the side chains of the complex to avoid steric clashes and to optimize atomic interactions with the new mutated residue. Since the template and mutant structures differ by only one mutation, the resulting model remains very close to the initial structure.

All wild-type (experimental) and mutant (modeled) structures can be downloaded at http://babylone.3bio.ulb.ac.be/DDGb_bias_structures/.

### 2.5 Prediction methods tested

We benchmarked the eight best known, available, and widely used ΔΔ*G*_*b*_ predictors published in recent years. We briefly describe their characteristics.

**mCSM-PPI2** [16] is a machine learning predictor that uses graph-based structural signatures of the inter-residue interaction network, evolutionary information, complex network metrics and energy terms.

**MutaBind2** [17] uses seven features including protein-solvent interactions, evolutionary conservation and physics-based thermodynamic stability.

**BeAtMuSiC** [12] is our in-house predictor. It estimates the ΔΔ*G*_*b*_ as a linear combination of the stability changes upon mutations (ΔΔ*G*) of the protein complex and of the individual interactants, computed by the PoPMuSiC predictor [45]. It uses statistical energy functions for ΔΔ*G* estimation, derived from the Boltzmann law which relates the frequency of occurrence of a structural pattern to its free energy.

**SSIPe** [18] combines protein interface profiles obtained from structure and sequence homology searches with physics-based energy functions.

**SAAMBE-3D** [19] is a machine learning-based predictor that utilizes 33 knowledge-based features representing the physical environment surrounding the mutation site.

**NetTree** [20] is a deep learning method based on convolutional neural networks and algebraic topology features. It uses element- and site-specific persistent homology to represent the structure of a protein complex and to translate it into topological features.

**FoldX** [46] is a purely physics-based method that uses empirical energy functions to predict ΔΔ*G*_*b*_ as described in the FOLDEF paper [10]. Its energy terms are defined by theoretical models (e.g. the van der Waals potential energy function), which are parameterized and weighted using empirical data.

**BindProfX** [15] combines the FoldX prediction and a profile score based on structural interface alignments obtained by the iAlign software [47]. The profile score exploits evolutionary information by comparing the frequencies of occurrence of the wild-type and the mutant amino acids in structurally similar interfaces. BindProfX is only applicable to protein dimers; when applied to higher-order multimers, we use the FoldX term only.

These predictors can be classified into three groups based on the nature of their approach: mCSM-PPI2, MutaBind2, SAAMBE-3D and NetTree are machine learning predictors whose features are extracted from protein structures, physics and evolution; SSIPe and BindProfX linearly combine an evolutionary term and a physics-based energy term using ΔΔ*G*_*b*_ data to optimize their models; BeAtMuSiC and FoldX are pure physics-based predictors.

In terms of training set, we have the following classification: NetTree was trained on antigen-antibody interaction data from the AB-Bind dataset [29] which is partially included in the SKEMPI 2.0 dataset; FoldX was trained on ΔΔ*G* data from ProTherm [23], however note that it has been updated several times since its first publication [10] in 2002 and it is unclear whether or not the current version (v5) [48] has used ΔΔ*G*_*b*_ data for parameterization; BeAtMuSiC was also trained on ΔΔ*G* values, with only two parameters to balance interprotein and intraprotein contributions adjusted using SKEMPI 1.0 ΔΔ*G*_*b*_ values; BindProfX was trained on SKEMPI 1.0 entries; all other predictors were trained on SKEMPI 2.0. Finally, mCSM-PPI2 and MutaBind2 included reverse mutations in addition to direct mutations in their training datasets.

Predictions from BeAtMuSiC, SSIPe, SAAMBE-3D, NetTree, BindProfX and FoldX were obtained by running their standalone code, while predictions from mCSM-PPI2 and MutaBind2 were obtained using their online webserver.

## 3 Results and discussion

### 3.1 An upper bound to the accuracy of predictors

Binding affinity change values collected from the literature and available in S2536 are derived from experiments performed using different techniques and under different environmental conditions such as pH, temperature or solvent additives. These differences add to the experimental error and usually lead to different ΔΔ*G*_*b*_ values for the same mutation in the same protein complex. Furthermore, although SKEMPI 2.0 is particularly well curated, curation errors cannot be avoided, as illustrated by the error corrections between SKEMPI 1.0 and SKEMPI 2.0 (see Supplementary Section 1). The uncertainty on ΔΔ*G*_*b*_ values places an upper bound on the precision of the predictions, which cannot exceed the accuracy of the experimental data.

An analytical method for estimating the upper bound on the Pearson correlation coefficient (*ρ*), which measures the strength of the linear relation between predicted and target values, and the lower bound on the root mean squared error (RMSE), which is a measure of the average error of a prediction, has recently been proposed [49, 50]. These bounds are expressed as:

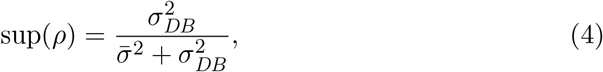

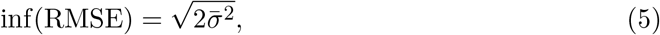

where 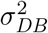 is the variance of ΔΔ*G*_*b*_ values in the whole dataset and 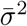 is the mean of the individual variances for redundant entries. We estimated the values of these bounds using the 116 redundant clusters with at least three entries among all single mutations from the SKEMPI 2.0 dataset.

We obtained: sup(*ρ*) = 0.89 and inf(RMSE) = 0.70 kcal/mol. Note, however, that these bounds are probably overestimated and underestimated, respectively, due to an underestimation of 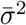. Indeed, only independent, uncorrelated, ΔΔ*G*_*b*_ measures of a given mutation can yield a correct estimation of the variance, which seems to not be always the case.

The performances of the tested predictors presented in the following sections can be compared with these “optimal” values. It should be stressed that an accuracy better than these bounds suggests that the predictor is overfitted towards the dataset. A good prediction should thus have a Pearson correlation significantly above zero but below the upper bound of 0.89. It is also expected to have an RMSE value above the lower bound of 0.70 kcal/mol. To give the reader an intuitive idea of the scale of the RMSE, we note that a predictor that consistently predicts ΔΔ*G*_*b*_ to be zero would obtain RMSE values of 2.3 and 1.8 kcal/mol on S2536 and C380, respectively.

### 3.2 Biases in the S2536 dataset

As mentioned by the SKEMPI authors [24, 25], mutations characterized and reported in the literature are not systematic but reflect the interests of the experimenters. The collected data have therefore biases towards specific residues, mutation types, spatial locations, proteins and protein families. These biases can lead to overoptimistic assessments of the predictors, even when strict cross-validation methods are used. Indeed, if training and test sets are subject to the same biases, a predictor can learn and replicate them, increasing both its apparent performance and generalization error. This can lead to a gap between the performances estimated from either a biased test set or a set of systematic mutations, raising concerns about the reliability of predictors. In this section we have quantified and discussed some of the biases in the S2536 mutations set.

First we note the imbalance in terms of mutation types. The occurrences of the 380 possible mutation types in S2536 are shown in Fig. 1a. Half of the mutations are towards alanine, 222 mutation types occur less than 5 times and 92 mutation types are not represented. This tendency is related to the prevalence of experimental alanine-scanning data in S2536. It may weaken the predictions of underrepresented mutation types.

**Figure 1:**
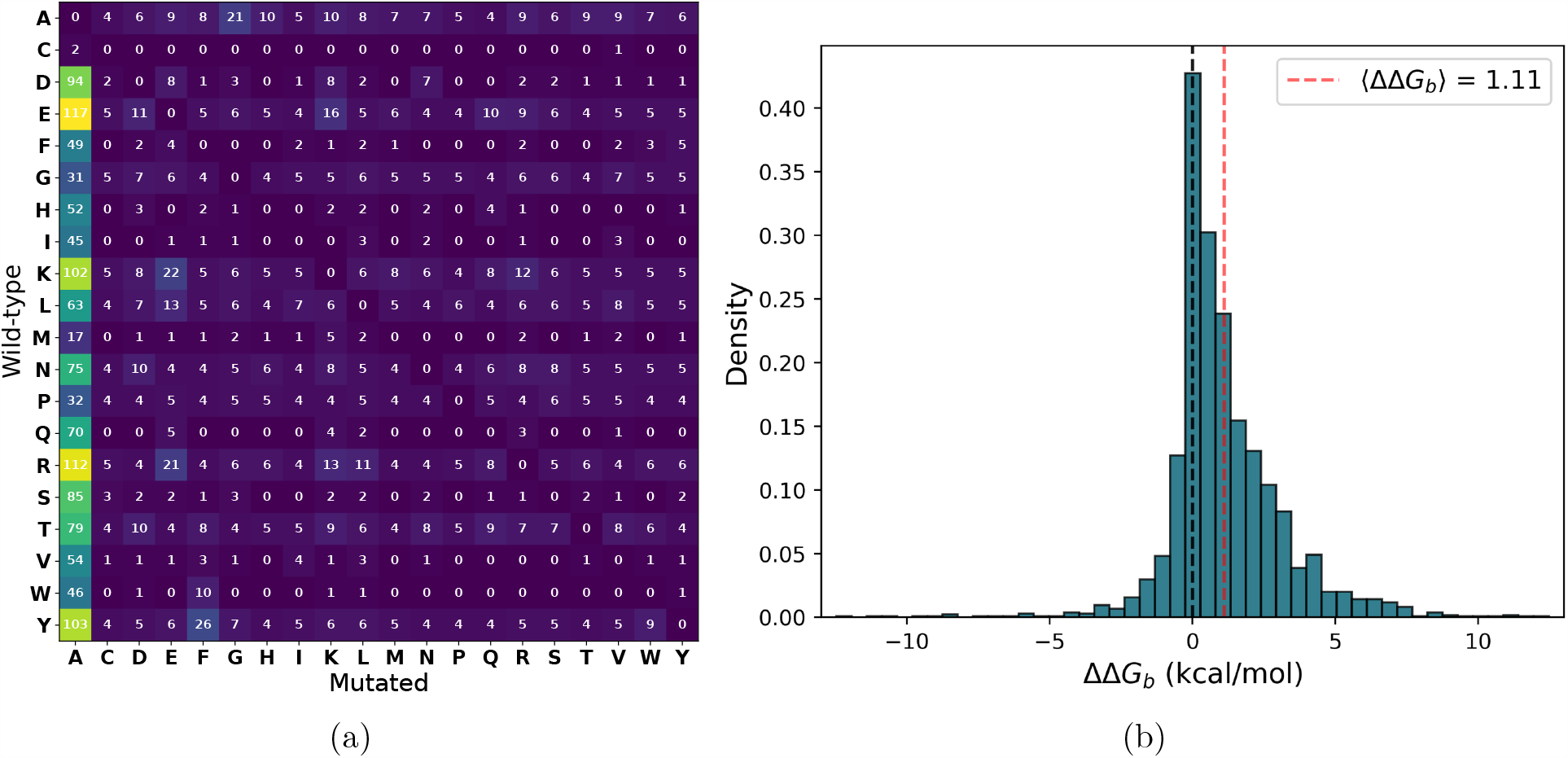
Characteristics of the S2536 dataset. (a) Number of occurrences of mutation types; (b) Distribution of the experimental ΔΔ*G*_*b*_ values (in kcal/mol).

Another notable imbalance is towards mutations located at protein-protein interfaces: 78% of S2536 entries are mutations of the 9% of residues located at the interface. Although though interface residues are usually more critical for the interaction, non-interface regions can also be important and their effects risk being overlooked by the predictors.

Finally, the ΔΔ*G*_*b*_ distribution is largely shifted towards positive values, as shown in Figure 1b. It has a mean value of 1.11 kcal/mol and a standard deviation of 1.99 kcal/mol with clear prevalence of destabilizing mutations. This imbalance is not surprising as experimentally studied complexes are often optimized for high binding affinity by evolution. However, it tends to cause predictors to systematically output destabilizing ΔΔ*G*_*b*_ values even for neutral and stabilizing mutations, thus preventing the symmetry property (Eq. (3)) from being satisfied. This issue, which is particularly problematic for, e.g., rational protein design, has been identified and widely investigated in the context of stability changes upon mutations [35, 33, 51, 52, 53, 34]. In the next sections, we will examine this in the context of changes in binding affinity.

Note that these imbalances were observed in S2536, but also occur in all single-site mutations of the SKEMPI 2.0 dataset (see Supplementary Section 2).

### 3.3 Performances on SKEMPI 2.0

We tested the performances of the eight selected predictors described in Methods (Section 2) on the direct and reverse mutations of the S2536 benchmark dataset. For that purpose, we used the Pearson correlation coefficient between predicted and experimental ΔΔ*G*_*b*_ values (*ρ*) as performance metric. The results are represented in Figures 2-3 and Table 1. Other metrics such as the root mean squared error (RMSE) and the Spearman rank correlation (*r*) lead to the similar conclusions (as shown in Table 1 and https://github.com/3BioCompBio/DDGb_bias).

**Table 1:** Performances of the eight benchmarked predictors measured by the Pearson correlation (*ρ*), the Spearman rank correlation (*r*) and root mean squared error (RMSE) on the datasets S2536-D, S2536-R, C380-D and C380-R.

| Predictors | S2536-D |  |  | S2536-R |  |  | C380-D |  |  | C380-R |  |  |
| --- | --- | --- | --- | --- | --- | --- | --- | --- | --- | --- | --- | --- |
| | $\rho$ | $r$ | RMSE | $\rho$ | $r$ | RMSE | $\rho$ | $r$ | RMSE | $\rho$ | $r$ | RMSE |
| mCSM-PPI2 | 0.91 | 0.90 | 0.96 | 0.78 | 0.71 | 1.37 | 0.45 | 0.48 | 1.27 | 0.39 | 0.36 | 1.59 |
| MutaBind2 | 0.90 | 0.85 | 0.92 | 0.59 | 0.47 | 1.62 | 0.47 | 0.58 | 1.20 | 0.38 | 0.39 | 1.40 |
| BeAtMuSiC | 0.34 | 0.40 | 1.90 | 0.09 | 0.03 | 2.66 | 0.35 | 0.34 | 1.26 | -0.05 | -0.02 | 2.37 |
| SSIPe | 0.53 | 0.46 | 1.76 | 0.47 | 0.35 | 2.16 | 0.37 | 0.41 | 1.30 | 0.23 | 0.23 | 2.05 |
| SAAMBE-3D | 0.88 | 0.85 | 1.02 | 0.11 | -0.05 | 2.53 | 0.16 | 0.14 | 1.31 | -0.17 | -0.08 | 2.45 |
| NetTree | 0.16 | 0.27 | 2.37 | -0.09 | -0.11 | 4.18 | 0.24 | 0.18 | 1.94 | -0.20 | -0.15 | 3.93 |
| BindProfX | 0.58 | 0.50 | 1.64 | 0.51 | 0.38 | 1.89 | 0.57 | 0.63 | 1.06 | 0.37 | 0.36 | 1.84 |
| FoldX | 0.44 | 0.48 | 1.99 | 0.31 | 0.34 | 2.20 | 0.33 | 0.55 | 2.32 | 0.14 | 0.36 | 2.31 |

**Figure 2:**
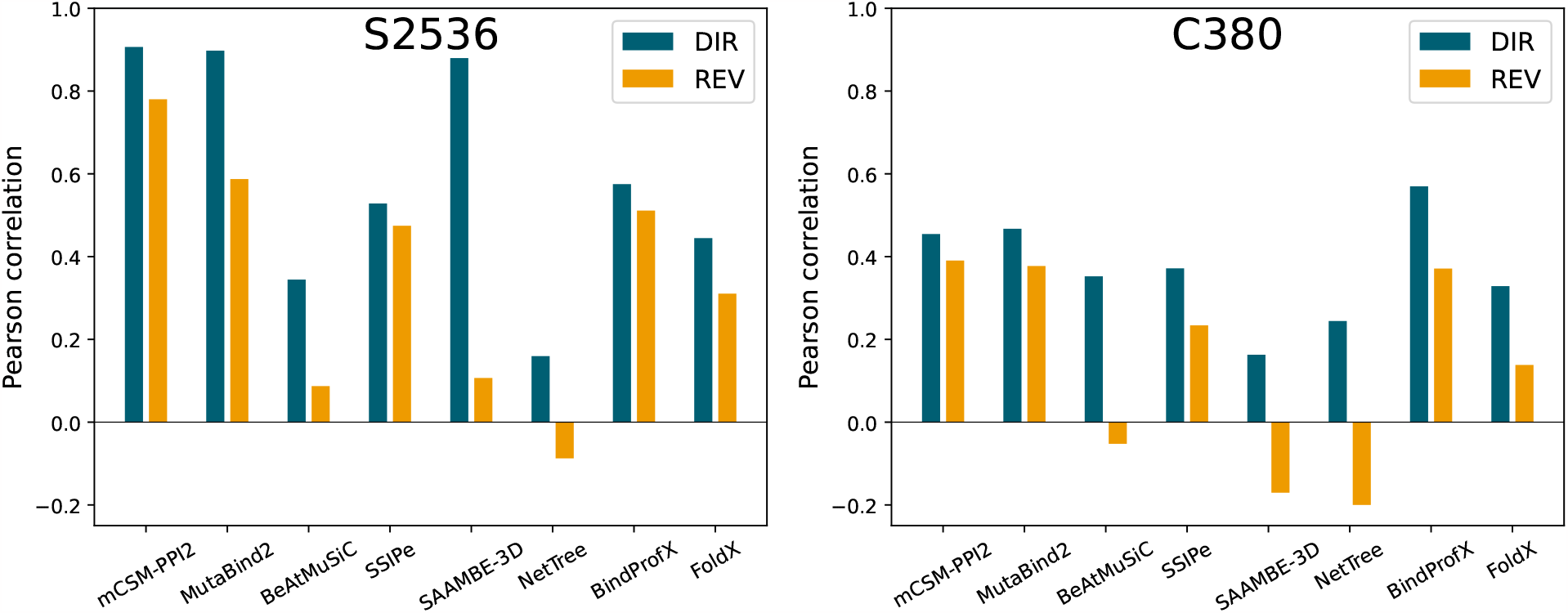
Pearson correlations *ρ* between experimental and predicted ΔΔ*G*_*b*_ values on direct (in blue) and reverse (in orange) mutations of S2536 (left) and C380 (right).

**Figure 3:**
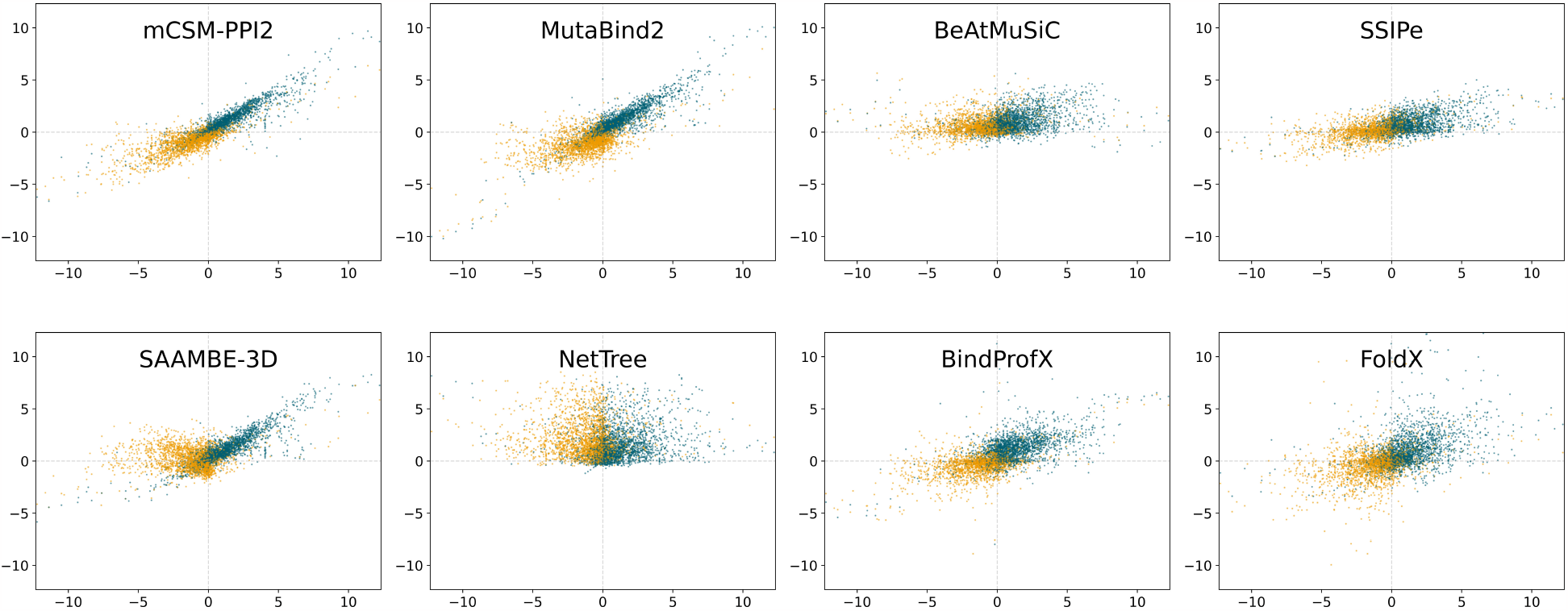
Predicted ΔΔ*G*_*b*_ values as a function of experimental ΔΔ*G*_*b*_ values (in kcal/mol) for the datasets S2536-D (blue dots) and S2536-R (orange dots). Predictions are obtained with mCSM-PPI2, MutaBind2, BeAtMuSiC, SSIPe, SAAMBE-3D, NetTree, BindProfX and FoldX.

This benchmark, though informative, should be considered with caution, as the extent of cross-validation differs according to the predictor. The main issue is that each of the benchmarked predictors is trained on a different subset of S2536, with various covering ratios (CR) with respect to the subset of direct (S2536-D) and reverse (S2536-R) mutations (Table 2). For instance, the training set of mCSM-PPI2 contains 99% of the S2536-D mutations, while that of NetTree only 10%. Furthermore, mCSM-PPI2 is trained on almost all reverse mutations of S2536-R and MutaBind2, on the fraction necessary to balance the number of stabilizing and destabilizing mutations.

**Table 2:**
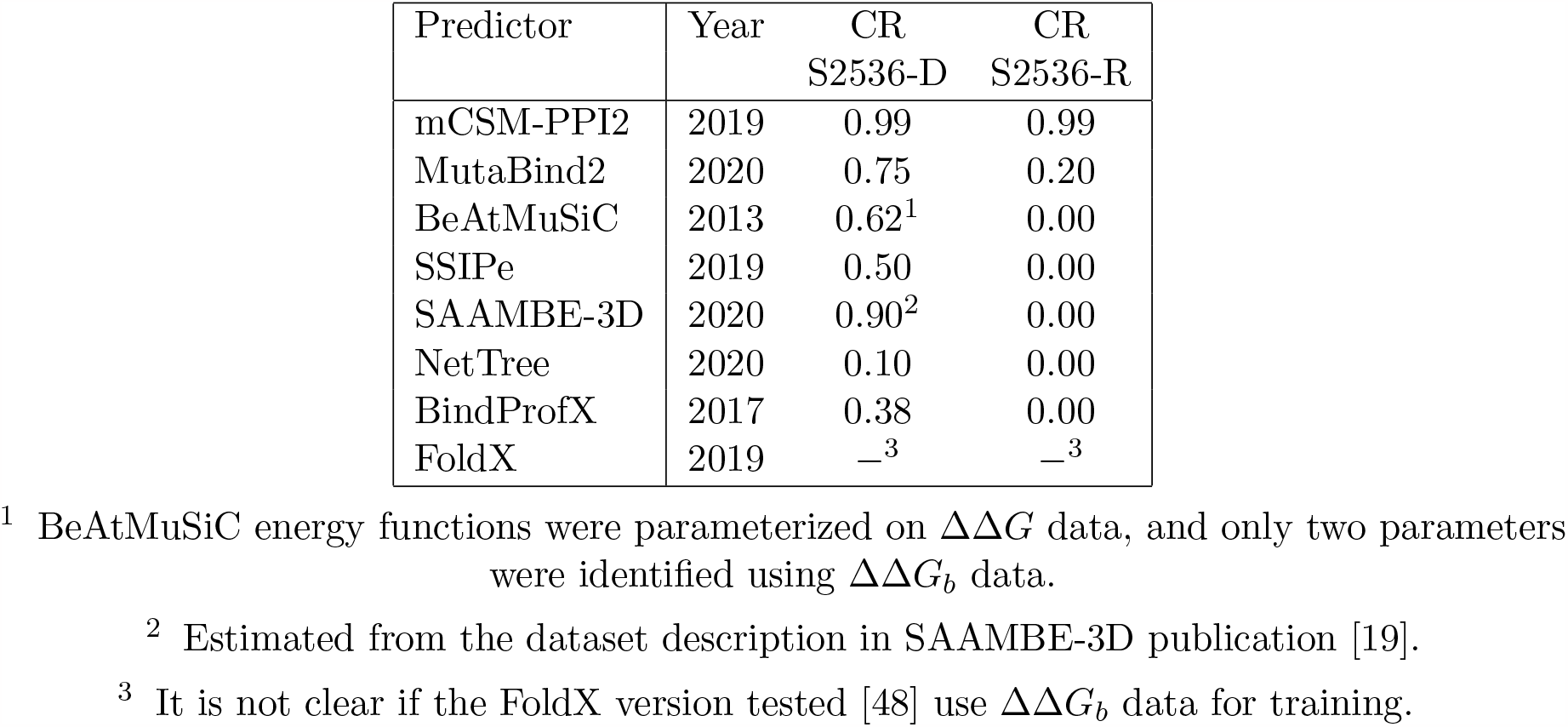
Year of publication of the eight benchmarked predictors and covering ratio (CR) of their training sets with respect to S2536-D and S2536-R.

The best performing predictors on the direct mutation set S2536-D are mCSM-PPI2, MutaBind2 and SAAMBE-3D with Pearson correlations *ρ* of 0.91, 0.90 and 0.88, respectively. These values exceed or are very close to the upper bound of 0.89 (Eq. (4)), which suggests some overfitting towards the training set. They are followed by BindProfX, SSIPe, FoldX, BeAtMuSiC and NetTree.

We observe that the performance of all predictors but SSIPe and BindProfX significantly drops when tested on reverse S2536-R mutations. The magnitude of the drop indicates how much each predictor is biased towards direct mutations, which are mostly destabilizing. mCSM-PPI2 and MutaBind2 perform the best on S2536-R, which is expected since they have reverse mutations in their training set; the performance of mCSM-PPI2 drops less than that of MutaBind2, probably because the latter have seen only a part of the reverse mutations during training.

Surprisingly, SSIPe and BindProfX are the most robust towards reverse mutations, with almost no drop in performance, although they do not use reverse mutations in training; their robustness is therefore not acquired by training but rather by the symmetry properties of the model. In contrast, BeAtMuSiC, SAAMBE-3D and NetTree basically fail to predict the ΔΔ*G*_*b*_ of reverse mutations. Note the particularly huge drop in performance of SAAMBE-3D, whose Pearson correlation decreases from 0.88 to 0.11; this predictor appears thus to be heavily biased towards destabilizing mutations.

This first benchmark shows that a bias toward destabilizing mutations is present in the context of ΔΔ*G*_*b*_ predictions. Note that the drop in performance observed when passing from direct to reverse mutations can partly be attributed to this bias but also to the presence of a larger proportion of mutations in S2536-R than in S2536-D which are unseen during training.

For the six methods trained on ΔΔ*G*_*b*_ data (mCSM-PPI2, MutaBind2, SSIPe, SAAMBE-3D, NetTree and BindProfX), the covering ratio CR between training and benchmark datasets accurately predicts the performances of the predictors. Indeed, we found an almost linear relationship between the CR of the six predictors and their Pearson correlation *ρ* on the S2536-D set, with a coefficient of determination *R*^2^ as high as 0.91 (Figure 4).

**Figure 4:**
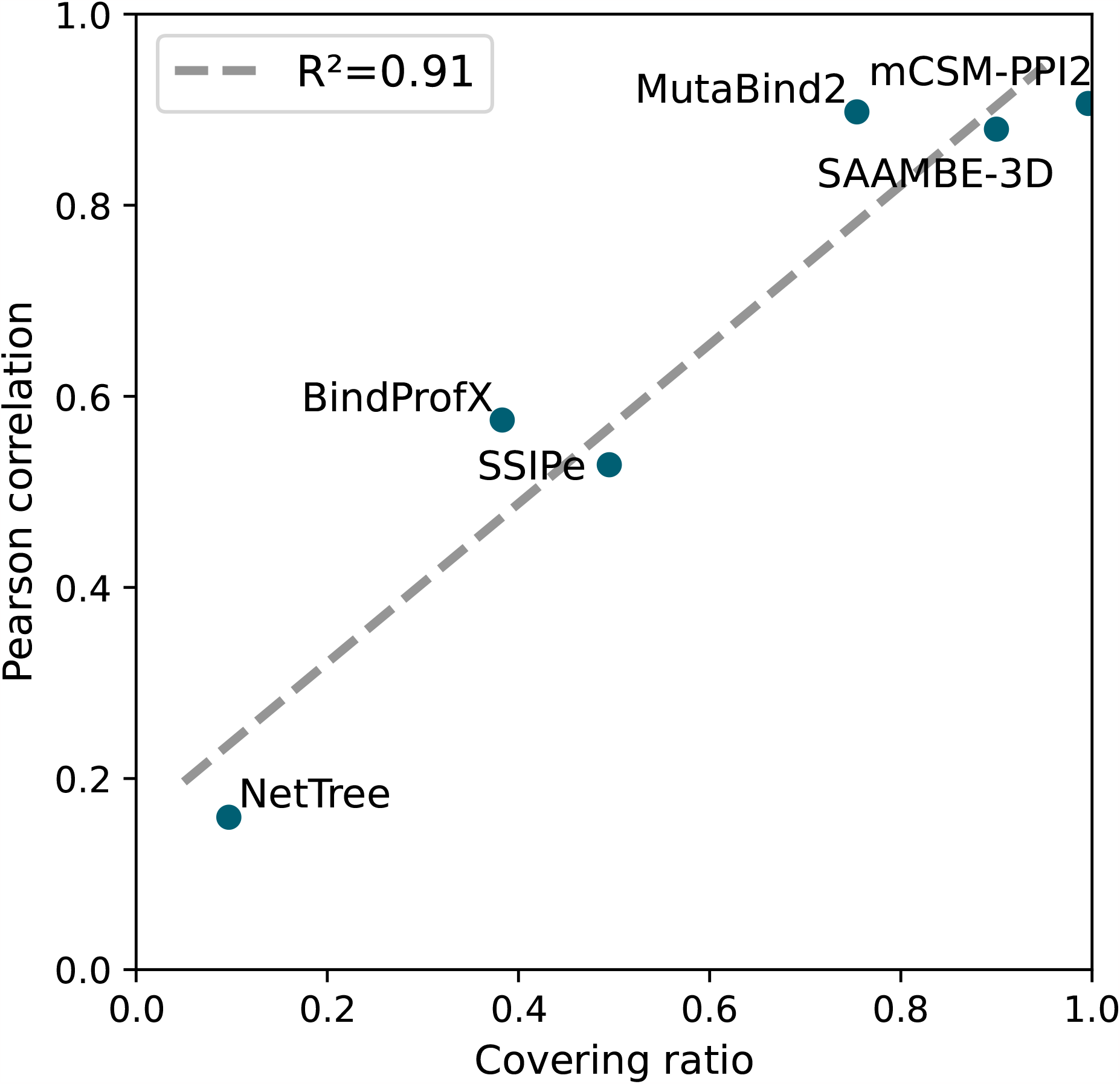
Relation between the covering ratio CR and the Pearson correlation *ρ* between predicted and experimental ΔΔ*G*_*b*_ values on the S2536-D set for six benchmarked predictors. The linear regression line (dashed) and coefficient of determination (*R*^2^) are indicated.

While this observation does not prove that these predictors are dataset specific and overfitted, it raises some concerns about their ability to generalize to mutations outside the training set. Therefore, further investigation based on a dataset of more systematic and unseen mutations is required: this is the topic of the next subsection.

### 3.4 Performances on SARS-CoV-2 mutations

The C380 dataset has two major advantages over S2536: it is unknown to the eight benchmarked predictors and it is systematic in terms of mutation types. This makes it a better dataset to evaluate the performances of the predictors. By comparing performances on direct and reverse mutations from C380-D and C380-R, we further explored the predictors’ bias towards destabilizing mutations; by comparing performances on mutations from S2536 and C380, we estimated the dataset dependence of the predictors. Predicted values and performance metrics on C380 are available on https://github.com/3BioCompBio/DDGb bias and predictions are graphically represented in Supplementary Figure S-4.

As shown in Figure 2, the performances of all predictors but NetTree drop from S2536 to C380, with no score higher than 0.6 on C380-D and 0.4 on C380-R. Comparing performances on direct mutations from the two datasets illustrates the heavy impact of the training dataset on the prediction accuracy, especially for the best performing predictors on the S2536 benchmark.

Note that the pure physics-based predictors (FoldX and BeAtMuSiC) and the predictors that use ΔΔ*G*_*b*_ data only to set up some weights and parameterize their model (BindProfX and SSIPe) only show relatively small drops in performance between S2536-D and C380-D. Among predictors which use machine learning more extensively, mCSM-PPI2 and MutaBind2 still show good performances on C380-D, ranking as second (MutaBind2: *ρ* = 0.47) and third (mCSM-PPI2: *ρ* = 0.45) after BindProfX (*ρ* = 0.57); their performance is, however, substantially reduced in comparison with the S2536-D benchmark; and SAAMBE-3D undergoes the largest performances drop.

The performance comparison between direct and reverse mutations of C380-D and C380-R confirms the conclusions of the previous section: all predictors suffer, to a different extent, from a bias towards destabilizing mutations. A way to quantify this bias for a given predictor is to compute the symmetry violation defined by Eq. (3) by computing the shift *δ* defined as:

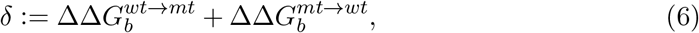

averaged over all C380 dataset entries. While some fluctuations in *δ* are expected and acceptable, a systematic deviation of the mean shift ⟨*δ*⟩ from zero quantifies the asymmetry of a predictor and its bias towards stabilizing or destabilizing mutations. A perfect unbiased value for ⟨*δ*⟩ is zero; its “worst-case” value can be estimated as twice the average ΔΔ*G*_*b*_ value in the dataset of direct mutations, which is 1.24 kcal/mol in C380. We thus estimated the “worst-case” *δ*-value to be about 2.5 kcal/mol.

We show in Figure 5 the distributions of *δ*-values for the eight predictors on C380. Analogous *δ*-values distributions are depicted for S2536 in Supplementary Figure S-5. We observe that all predictors have a statistically significant shift towards destabilizing mutations, with a vanishing *p*-value, but amplitude of the shift widely varies. The most symmetric predictors are, as expected, those that perform best on reverse mutations: MutaBind2 with ⟨*δ*⟩ = 0.28 kcal/mol followed by mCSM-PPI2 with ⟨*δ*⟩ = 0.47 kcal/mol.

**Figure 5:**
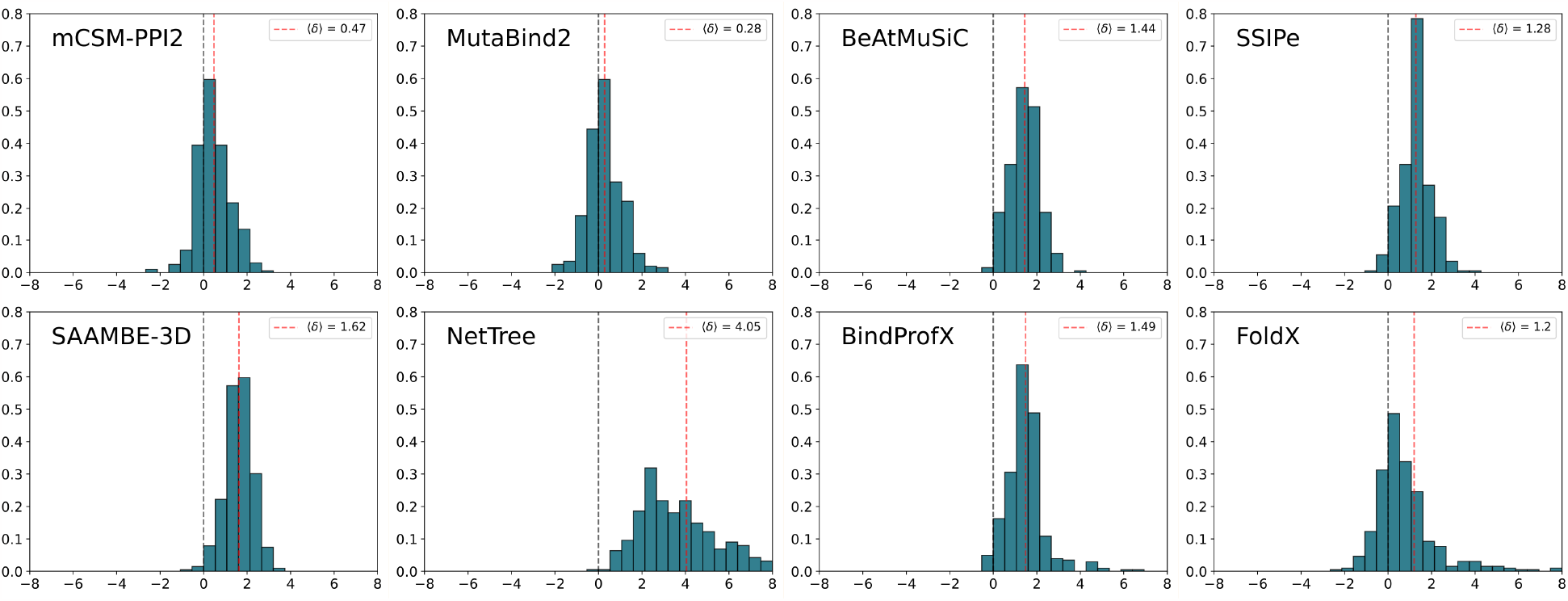
Distribution of the shift *δ* (in kcal/mol) for the eight benchmarked predictors calculated for mutations from C380. The vertical blue dashed lines indicate *δ* = 0 and the vertical red dashed lines, the value of ⟨*δ*⟩.

This confirms that the usage of reverse mutations for training can largely reduce the asymmetry of the predictions. More biased predictions are observed for FoldX, SSIPe, BeAtMuSiC, BindProfX and SAAMBE-3D, with ⟨*δ*⟩ = 1.20, 1.28, 1.44, 1.49 and 1.62 kcal/mol, respectively. These values indicate a bias towards destabilizing mutations, which is, however, still significantly lower than the “worst-case” bias. This means that such predictions are still able to distinguish the tendency between a set of mostly-stabilizing and mostly-destabilizing mutations. In contrast, NetTree obtains ⟨*δ*⟩ = 4.05 kcal/mol, which is largely above the “worst-case” bias and reflects its inability to distinguish stabilizing from destabilizing mutations. This particularly large ⟨*δ*⟩-value can partly be explained by NetTree’s tendency to predict very large ΔΔ*G*_*b*_ values of about 2 kcal/mol, much higher than average experimental values.

In summary, this benchmark represents a fair and objective way to evaluate the performance of the predictors, since C380 is unknown to all. It confirms the presence of biases towards destabilizing mutations in the state-of-the-art ΔΔ*G*_*b*_ predictors and highlights the two predictors mCSM-PPI2 and MutaBind2 that are the least affected by this bias.

### 3.5 Performances and biases towards mutation properties

We investigated the predictors’ performances on subsets of S2536-D containing mutations sharing similar properties, i.e., mutation type, mutation location and type of complex, in order to highlight the predictors’ strengths and weaknesses. As the standard deviations *σ* of the experimental ΔΔ*G*_*b*_ values widely differ according to the subset, we used the normalized RMSE defined as nRMSE := RMSE*/σ* to assess the predictions. The results are shown in Figure 6. All observations discussed below are statistically significant with almost vanishing *p*-values (*<* 0.0001).

**Figure 6:**
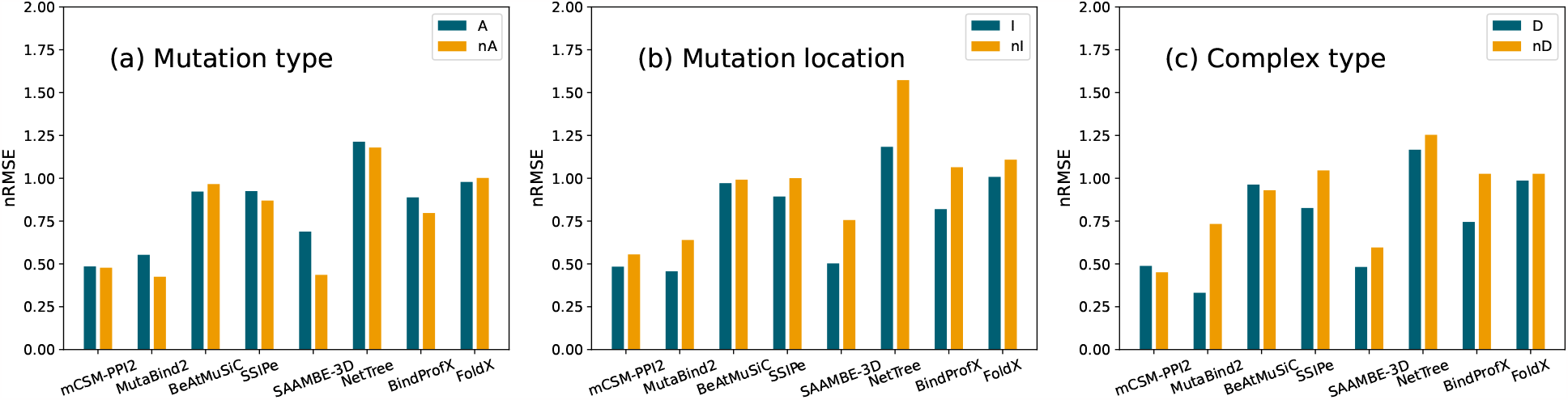
Normalized root mean squared error (nRMSE) of the eight predictors on subsets of S2536-D. Subsets were defined based on (a) mutation type: mutation towards Ala (A) vs other mutations (nA); (b) mutation location: mutations at the interface (I) vs other mutations (nI). (c) complex type: mutation on dimeric complexes (D) vs mutations on multi-n-meric complexes (*n >* 2) (nD).

We first analyzed separately the subset of mutations towards alanine and the subset of other mutations. As seen in Figure 6a, no substantial differences are observed between these two subsets, except that MutaBind2 and SAAMBE-3D perform slightly better on the latter subset. This might be explained by actual strengths/weaknesses of the predictors or could suggest a mild overfitting, since it is easier to memorize ΔΔ*G*_*b*_ values on underrepresented mutation types.

Most predictors are slightly weaker on mutations outside the protein-protein interface (Figure 6b). This is foreseeable, since effects on binding affinity of non-interface mutations are indirect an thus more difficult to predict. MutaBind2, BindProfX, SAAMBE-3D and NetTree suffer from the largest increase in nRMSE. In contrast, BeAtMuSiC and FoldX present similar performances on both subsets. SSIPe shows a surprisingly small drop in performance on mutations outside the interface, although it explicitly claims to be only able to predict interface mutations.

When comparing mutations in dimers to mutations in higher-order multimers (Figure 6c), we observe that mCSM-PPI2, BeAtMuSiC and FoldX are the most stable and that MutaBind2, SSIPe, SAAMBE-3D and BindProfX show the largest performance drop. SSIPe’s poor performance on higher-order multimers is not surprising as it explicitly announces not to predict such mutations. BindProfX’s drop is related to the fact that its predictions on higher-order multimers are taken from FoldX (see Methods). Paradoxically, mCSM-PPI2 does not require specifying which chains make up the two interactants, although higher-order multimers have several protein-protein interfaces and so there is an ambiguity. In spite of this, it maintains the same performance on both subsets, which could suggest overfitting towards its training dataset. In contrast, MutaBind2 asks the chains included in the interactants, but has the largest performance drop on higher-order multimers.

We also assessed the performances on other S2536-D subsets, partitioned by secondary structure, solvent exposure in the complex and interface sub-regions [54] (definitions in Supplementary Section 4), but no relevant observations where found. Results are available at https://github.com/3BioCompBio/DDGb_bias).

### 3.6 Strategies for avoiding biased predictions

To ensure the generalizability of the predictions, *k*-fold cross-validation procedures should be carefully performed, avoiding blindly splitting the training set. Indeed, when separating a dataset into folds, a direct mutation and its corresponding reverse mutation should end up in the same fold to avoid that information from one mutation influences the prediction of the other. As the S2536 dataset contains multiple homologous complexes differing by only a few mutations, random cross-validations can also lead to information leaks from training to testing sets and provide overoptimistic results. Thus, mutations on homologous complexes should also be kept in the same fold [24].

However, dataset biases can be learned by the predictors even if a strict cross-validation procedure is used. To illustrate this, we started by noticing that half of the mutations from S2536-D are towards alanine (*X* → *A*) and thus that half of the mutations from S2536-R are from alanine (*A* → *X*). Knowing moreover that S2536-D and S2536-R contain mostly destabilizing and mostly stabilizing mutations, respectively, the sign of ΔΔ*G*_*b*_ can be often correctly guessed for *X* → *A* and *A* → *X* mutations while holding no predictive power. In other words, predictors can learn imbalances and cross correlations between mutations’ properties from S2536, which improves its performances in cross-validation while also increasing its generalization error.

As a proof of this phenomenon, we created a “perfectly biased” predictor, which estimates ΔΔ*G*_*b*_ as the mean of the experimental ΔΔ*G*_*b*_ values of the same mutation type in the training set (or zero if the mutation type was never encountered). This predictor manages to obtain a Pearson correlation *ρ* = 0.46 on S2536-DR in 10-fold cross-validation. When applying the same predictor (trained on S2536-DR) on mutation type-balanced, interface-only entries from C380-DR, the Pearson correlation falls to *ρ* = 0.35, and completely vanishes when dropping the interface filter and applying the predictor to the whole dataset of mutations on the RBD-ACE2 complex (-DR) with *ρ* = 0.04. The same phenomenon also happens, with however slightly smaller correlations, when considering direct mutations only. We indeed found *ρ* = 0.34 in 10-fold cross-validation on S2536-D, *ρ* = 0.27 on C380-D and *ρ* = 0.05 on RBD-ACE2 (-D). Note that these scores are only an underestimation of how dataset-dependent cross correlations from S2536 can impact predictions; we have indeed only considered mutation type-related biases.

As extensively discussed above, asymmetric predictions are another type of unwanted bias. One easy way to avoid it is to symmetrize the prediction results. Indeed, the prediction shift *δ* vanishes when redefining the prediction of a mutation *wt* → *mt* as:

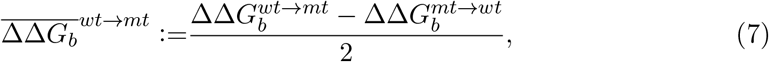

with, as a consequence, 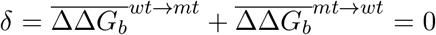. This operation requires both wild-type and mutant structures, but does not introduce any internal modifications to the predictor itself. Some but not all mutant structures have been resolved experimentally; we listed in the https://github.com/3BioCompBio/DDGb bias repository the pairs of resolved wild-type and mutant structures from SKEMPI 2.0 that are separated by a single mutation (more details in Supplementary Section 5). Alternatively, the unavailable mutant structures can be modeled with homology modeling techniques using the wild-type structure as a template.

Symmetrized versions of all tested predictors were obtained using Eq. (7). For predictors that suffer from a strong bias towards destabilizing mutations, the Pearson correlation coefficient of the symmetrized version falls somewhere between their scores on direct and on reverse mutations. In contrast, the least asymmetric predictors, mCSM-PPI2, MutaBind2, BindProfX, FoldX and SSIPe, show a significantly improved score on the reverse datasets S2536-R and C380-R, as well on the combined datasets S2536-DR and C380-DR, and similar or only slightly lower performance on the direct datasets S2536-D and C380-D (Supplementary Section 3). This shows that the overall performances of some predictors can be improved while also increasing their symmetry without introducing any internal changes to the model.

As seen in the previous subsections, an alternative strategy to reduce the asymmetry of the predictions consists in using reverse mutations for training. Among the tested predictors, MutaBind2 and mCSM-PPI2 apply this technique and reach good symmetry properties. This practice increases the generalizability and robustness of predictors. However, the symmetrization of the training set has to be done carefully.

Indeed, due to the presence of wild-type/mutant pairs in SKEMPI 2.0, adding the reverse of all mutations, as done in mCSM-PPI2, leads to redundant entries that should be avoided, as they are a source of biases.

### 3.7 Predictors’ computational efficiency

Computational time efficiency is another characteristic to consider when choosing a prediction method, especially when a large set of mutations has to be analyzed, as for example in the study of variants impact on the interactome [2]. In terms of speed, BeAtMuSiC and SAAMBE-3D are fast enough to enable large-scale computational mutagenesis experiments; indeed, they are able to predict all possible single-site mutations in a protein complex in a few to a few tens of seconds. While FoldX is significantly slower, it still can perform all mutations in a small protein complex in about a few hours. In contrast, mCSM-PPI2, MutaBind2, SSIPe, NetTree and BindProfX are time consuming and require tens of seconds to tens of minutes to run a single mutation. This prevents their use for large-scale applications.

## 4 Conclusions

In the last decade, the computational prediction of how mutations impact protein-protein binding affinity have experienced substantial improvements. Thanks to the large amount of experimental mutagenesis data generated and the development of new machine learning algorithms and accurate force fields, many ΔΔ*G*_*b*_ predictors that reach good performance have been developed and used in biotechnological and biopharmaceutical applications.

However, as clearly illustrated in our benchmarking analyses, the predictive power of a method is not necessarily well represented by its scores on its training dataset even if a strict cross-validation procedure is used. This makes the validation process particularly challenging. Here we identified two main issues, which are the predictors’ systematic asymmetry and their lack of generalization on mutations outside their training set. They are discussed below.

### Lack of generalization

A major challenge in ΔΔ*G*_*b*_ predictions is to distinguish between statistical relations that are dataset dependent and the “true” ones that have a biological meaning. We would like to stress that, while physics- and evolution-based methods are at least partly equipped to tackle this problem, pure machine learning methods struggle to make this distinction. This can explain the particularly large performance drop on unknown mutations observed for most purely machine learning methods such as SAAMBE-3D and the good generalizability properties observed for methods that are totally or partly physics-based, such as BeAtMuSiC, BindProfX and FoldX.

The generalizability of a predictor must be tested on independent sets of mutations outside the training set. Sets of systematic mutations obtained by deep mutagenesis experiments, such as C380, have the advantage of not being impacted by literature biases. They are thus appropriate for validating and benchmarking predictions, even though their ΔΔ*G*_*b*_ values are less accurate than those obtained by individual thermodynamic experiments.

### Symmetry properties

Symmetry properties should be carefully checked when constructing a prediction model. One way to assess them is on the basis of the shift *δ* (Eq. (6)). As a general rule, the symmetry of a predictor can be achieved by: (1) using symmetric data during training by including all or a fraction of reverse mutations, as done in mCSM-PPI2 and MutaBind2; (2) enforcing symmetry in the predictor’s mathematical model, as in [33]; and (3) applying symmetry-correction methods through, e.g., the symmetrization defined in Eq. (7). Method (1) is a good practice which, as we showed, can increase the generalizability of the predictions. Method (2) can help the predictor to be symmetric, but it is only applicable when the mathematical expression of the model is known. Method (3) is the easiest to implement, but is efficient only if the predictor is already robust to symmetry.

There are additional challenges that need to be addressed. First, further data on binding affinity and interactions need to be collected. Accurate ΔΔ*G*_*b*_ thermodynamics data have not been systematically collected for the past five years, after SKEMPI 2.0’s release. Also, deep mutagenesis data of binding affinity are currently generated at a high rate but need to be collected, curated and harmonized. Secondly, the interpretation of ΔΔ*G*_*b*_ prediction models is an issue that we do not explore in this paper and that is not sufficiently discussed in the literature. Indeed, performance is not the only criterion for evaluating a prediction model. Insights into model interpretation can help gaining physical understanding of molecular recognition and protein-protein binding mechanisms.

Finally, there is a need for more independent assessments. We invite the community to set up blind challenges for the prediction of changes in protein-protein binding affinity upon mutations, similar to what has been done during the 26^*th*^ critical assessment of predicted interactions (CAPRI) experiment [55]. These community-wide blind challenges provide important insights into whether and how different predictors achieve the targeted accuracy, and help drive the development of new methods.

## References

[1] Nidhi Sahni, Song Yi, Mikko Taipale, Juan I Fuxman Bass, Jasmin Coulombe-Huntington, Fan Yang, Jian Peng, Jochen Weile, Georgios I Karras, Yang Wang, et al. Widespread macromolecular interaction perturbations in human genetic disorders. Cell, 161(3):647–660, 2015.

[2] Feixiong Cheng, Junfei Zhao, Yang Wang, Weiqiang Lu, Zehui Liu, Yadi Zhou, William R Martin, Ruisheng Wang, Jin Huang, Tong Hao, et al. Comprehensive characterization of protein–protein interactions perturbed by disease mutations. Nature Genetics, 53(3):342–353, 2021.

[3] Anupama Yadav, Marc Vidal, and Katja Luck. Precision medicine—networks to the rescue. Current Opinion in Biotechnology, 63:177–189, 2020.

[4] Hongzhu Cui, Nan Zhao, and Dmitry Korkin. Multilayer view of pathogenic SNVs in human interactome through in silico edgetic profiling. Journal of Molecular Biology, 430(18):2974–2992, 2018.

[5] Laura Nevola and Ernest Giralt. Modulating protein–protein interactions: the potential of peptides. Chemical Communications, 51(16):3302–3315, 2015.

[6] Haiying Lu, Qiaodan Zhou, Jun He, Zhongliang Jiang, Cheng Peng, Rongsheng Tong, and Jianyou Shi. Recent advances in the development of protein–protein interactions modulators: mechanisms and clinical trials. Signal Transduction and Targeted Therapy, 5(1):1–23, 2020.

[7] Stephanie Leavitt and Ernesto Freire. Direct measurement of protein binding energetics by isothermal titration calorimetry. Current Opinion in Structural Biology, 11(5):560–566, 2001.

[8] Katja Luck, Dae-Kyum Kim, Luke Lambourne, Kerstin Spirohn, Bridget E Begg, Wenting Bian, Ruth Brignall, Tiziana Cafarelli, Francisco J Campos-Laborie, Benoit Charloteaux, et al. A reference map of the human binary protein interactome. Nature, 580(7803):402–408, 2020.

[9] Tanja Kortemme and David Baker. A simple physical model for binding energy hot spots in protein–protein complexes. Proceedings of the National Academy of Sciences, 99(22):14116–14121, 2002.

[10] Raphael Guerois, Jens Erik Nielsen, and Luis Serrano. Predicting changes in the stability of proteins and protein complexes: a study of more than 1000 mutations. Journal of Molecular Biology, 320(2):369–387, 2002.

[11] Song Liu, Chi Zhang, Hongyi Zhou, and Yaoqi Zhou. A physical reference state unifies the structure-derived potential of mean force for protein folding and binding. Proteins: Structure, Function, and Bioinformatics, 56(1):93–101, 2004.

[12] Yves Dehouck, Jean Marc Kwasigroch, Marianne Rooman, and Dimitri Gilis. BeAtMuSiC: prediction of changes in protein–protein binding affinity on mutations. Nucleic Acids Research, 41(W1):W333–W339, 2013.

[13] Douglas EV Pires, David B Ascher, and Tom L Blundell. mCSM: predicting the effects of mutations in proteins using graph-based signatures. Bioinformatics, 30(3):335–342, 2014.

[14] Minghui Li, Franco L Simonetti, Alexander Goncearenco, and Anna R Panchenko. MutaBind estimates and interprets the effects of sequence variants on protein– protein interactions. Nucleic Acids Research, 44(W1):W494–W501, 2016.

[15] Peng Xiong, Chengxin Zhang, Wei Zheng, and Yang Zhang. BindProfX: assessing mutation-induced binding affinity change by protein interface profiles with pseudo-counts. Journal of Molecular Biology, 429(3):426–434, 2017.

[16] Carlos HM Rodrigues, Yoochan Myung, Douglas EV Pires, and David B Ascher. mCSM-PPI2: predicting the effects of mutations on protein–protein interactions. Nucleic Acids Research, 47(W1):W338–W344, 2019.

[17] Ning Zhang, Yuting Chen, Haoyu Lu, Feiyang Zhao, Roberto Vera Alvarez, Alexander Goncearenco, Anna R Panchenko, and Minghui Li. MutaBind2: predicting the impacts of single and multiple mutations on protein-protein interactions. Iscience, 23(3):100939, 2020.

[18] Xiaoqiang Huang, Wei Zheng, Robin Pearce, and Yang Zhang. SSIPe: Accurately estimating protein–protein binding affinity change upon mutations using evolutionary profiles in combination with an optimized physical energy function. Bioinformatics, 36(8):2429–2437, 2020.

[19] Swagata Pahari, Gen Li, Adithya Krishna Murthy, Siqi Liang, Robert Fragoza, Haiyuan Yu, and Emil Alexov. SAAMBE-3D: predicting effect of mutations on protein–protein interactions. International Journal of Molecular Sciences, 21(7):2563, 2020.

[20] Menglun Wang, Zixuan Cang, and Guo-Wei Wei. A topology-based network tree for the prediction of protein–protein binding affinity changes following mutation. Nature Machine Intelligence, 2(2):116–123, 2020.

[21] Carlos HM Rodrigues, Douglas EV Pires, and David B Ascher. mmCSM-PPI: predicting the effects of multiple point mutations on protein–protein interactions. Nucleic Acids Research, 49(W1):W417–W424, 2021.

[22] Themis Lazaridis and Martin Karplus. Effective energy functions for protein structure prediction. Current Opinion in Structural Biology, 10(2):139–145, 2000.

[23] M Michael Gromiha, Jianghong An, Hidetoshi Kono, Motohisa Oobatake, Hatsuho Uedaira, Ponraj Prabakaran, and Akinori Sarai. ProTherm, version 2.0: thermodynamic database for proteins and mutants. Nucleic Acids Research, 28(1):283–285, 2000.

[24] Iain H Moal and Juan Fernández-Recio. SKEMPI: a structural kinetic and energetic database of mutant protein interactions and its use in empirical models. Bioinformatics, 28(20):2600–2607, 2012.

[25] Justina Jankauskaitė, Brian Jiménez-García, Justas Dapkūnas, Juan Fernández-Recio, and Iain H Moal. SKEMPI 2.0: an updated benchmark of changes in protein–protein binding energy, kinetics and thermodynamics upon mutation. Bioinformatics, 35(3):462–469, 2019.

[26] Kurt S Thorn and Andrew A Bogan. ASEdb: a database of alanine mutations and their effects on the free energy of binding in protein interactions. Bioinformatics, 17(3):284–285, 2001.

[27] MD Shaji Kumar and M Michael Gromiha. PINT: protein–protein interactions thermodynamic database. Nucleic Acids Research, 34(uppl 1):D195–D198, 2006.

[28] Panagiotis L Kastritis, Iain H Moal, Howook Hwang, Zhiping Weng, Paul A Bates, Alexandre MJJ Bonvin, and Jöel Janin. A structure-based benchmark for protein– protein binding affinity. Protein Science, 20(3):482–491, 2011.

[29] Sarah Sirin, James R Apgar, Eric M Bennett, and Amy E Keating. AB-Bind: antibody binding mutational database for computational affinity predictions. Protein Science, 25(2):393–409, 2016.

[30] Sherlyn Jemimah, K Yugandhar, and M Michael Gromiha. PROXiMATE: a database of mutant protein–protein complex thermodynamics and kinetics. Bioinformatics, 33(17):2787–2788, 2017.

[31] Quanya Liu, Peng Chen, Bing Wang, Jun Zhang, and Jinyan Li. dbMPIKT: a database of kinetic and thermodynamic mutant protein interactions. BMC Bioinformatics, 19(1):1–7, 2018.

[32] Thom Vreven, Iain H Moal, Anna Vangone, Brian G Pierce, Panagiotis L Kastritis, Mieczyslaw Torchala, Raphael Chaleil, Brian Jiménez-García, Paul A Bates, Juan Fernandez-Recio, et al. Updates to the integrated protein–protein interaction benchmarks: docking benchmark version 5 and affinity benchmark version 2. Journal of Molecular Biology, 427(19):3031–3041, 2015.

[33] Fabrizio Pucci, Katrien Bernaerts, Fabian Teheux, Dimitri Gilis, and Marianne Rooman. Symmetry principles in optimization problems: an application to protein stability prediction. IFAC-PapersOnLine, 48(1):458–463, 2015.

[34] Dinara R Usmanova, Natalya S Bogatyreva, Joan Ariño Bernad, Aleksandra A Eremina, Anastasiya A Gorshkova, German M Kanevskiy, Lyubov R Lonishin, Alexander V Meister, Alisa G Yakupova, Fyodor A Kondrashov, and Dmitry N Ivankov. Self-consistency test reveals systematic bias in programs for prediction change of stability upon mutation. Bioinformatics, 34(21):3653–3658, 2018.

[35] Fabrizio Pucci, Katrien V Bernaerts, Jean Marc Kwasigroch, and Marianne Rooman. Quantification of biases in predictions of protein stability changes upon mutations. Bioinformatics, 34(21):3659–3665, 2018.

[36] Tyler N Starr, Allison J Greaney, Sarah K Hilton, Daniel Ellis, Katharine HD Crawford, Adam S Dingens, Mary Jane Navarro, John E Bowen, M Alejandra Tortorici, Alexandra C Walls, et al. Deep mutational scanning of SARS-CoV-2 receptor binding domain reveals constraints on folding and ACE2 binding. Cell, 182(5):1295–1310, 2020.

[37] George D Rose, Ari R Geselowitz, Glenn J Lesser, Richard H Lee, and Micheal H Zehfus. Hydrophobicity of amino acid residues in globular proteins. Science, 229(4716):834–838, 1985.

[38] Georgios A Dalkas, Fabian Teheux, Jean Marc Kwasigroch, and Marianne Rooman. Cation–π, amino–π, π–π, and H-bond interactions stabilize antigen– antibody interfaces. Proteins: Structure, Function, and Bioinformatics, 82(9):1734–1746, 2014.

[39] Wolfgang Kabsch and Christian Sander. Dictionary of protein secondary structure: pattern recognition of hydrogen-bonded and geometrical features. Biopolymers: Original Research on Biomolecules, 22(12):2577–2637, 1983.

[40] Helen M Berman, John Westbrook, Zukang Feng, Gary Gilliland, Talapady N Bhat, Helge Weissig, Ilya N Shindyalov, and Philip E Bourne. The protein data bank. Nucleic Acids Research, 28(1):235–242, 2000.

[41] Stephen M Lu, Wuyuan Lu, MA Qasim, Stephen Anderson, Izydor Apostol, Wojciech Ardelt, Theresa Bigler, Yi Wen Chiang, James Cook, Michael NG James, et al. Predicting the reactivity of proteins from their sequence alone: Kazal family of protein inhibitors of serine proteinases. Proceedings of the National Academy of Sciences, 98(4):1410–1415, 2001.

[42] Henrik Gardsvoll, Bernard Gilquin, Marie Hélène Le Du, Andre M énèz, Thomas JD Jørgensen, and Michael Ploug. Characterization of the functional epitope on the urokinase receptor: complete alanine scanning mutagenesis supplemented by chemical cross-linking. Journal of Biological Chemistry, 281(28):19260–19272, 2006.

[43] Jun Lan, Jiwan Ge, Jinfang Yu, Sisi Shan, Huan Zhou, Shilong Fan, Qi Zhang, Xuanling Shi, Qisheng Wang, Linqi Zhang, et al. Structure of the SARS-CoV-2 spike receptor-binding domain bound to the ACE2 receptor. Nature, 581(7807):215–220, 2020.

[44] Benjamin Webb and Andrej Sali. Comparative protein structure modeling using MODELLER. Current Protocols in Bioinformatics, 54(1):5–6, 2016.

[45] Yves Dehouck, Aline Grosfils, Benjamin Folch, Dimitri Gilis, Philippe Bogaerts, and Marianne Rooman. Fast and accurate predictions of protein stability changes upon mutations using statistical potentials and neural networks: PoPMuSiC-2.0. Bioinformatics, 25(19):2537–2543, 2009.

[46] Joost Schymkowitz, Jesper Borg, Francois Stricher, Robby Nys, Frederic Rousseau, and Luis Serrano. The FoldX web server: an online force field. Nucleic Acids Research, 33(uppl 2):W382–W388, 2005.

[47] Mu Gao and Jeffrey Skolnick. iAlign: a method for the structural comparison of protein–protein interfaces. Bioinformatics, 26(18):2259–2265, 2010.

[48] Javier Delgado, Leandro G Radusky, Damiano Cianferoni, and Luis Serrano. FoldX 5.0: working with rna, small molecules and a new graphical interface. Bioinformatics, 35(20):4168–4169, 2019.

[49] Ludovica Montanucci, Pier Luigi Martelli, Nir Ben-Tal, and Piero Fariselli. A natural upper bound to the accuracy of predicting protein stability changes upon mutations. Bioinformatics, 35(9):1513–1517, 2019.

[50] Silvia Benevenuta and Piero Fariselli. On the upper bounds of the real-valued predictions. Bioinformatics and Biology Insights, 13:1177932219871263, 2019.

[51] Octav Caldararu, Rukmankesh Mehra, Tom L Blundell, and Kasper P Kepp. Systematic investigation of the data set dependency of protein stability predictors. Journal of Chemical Information and Modeling, 60(10):4772–4784, 2020.

[52] Kristoffer T Bæk and Kasper P Kepp. Data set and fitting dependencies when estimating protein mutant stability: Toward simple, balanced, and interpretable models. Journal of Computational Chemistry, 43(8):504–518, 2022.

[53] Vladimir Potapov, Mati Cohen, and Gideon Schreiber. Assessing computational methods for predicting protein stability upon mutation: good on average but not in the details. Protein Engineering, Design & Selection, 22(9):553–560, 2009.

[54] Emmanuel D Levy. A simple definition of structural regions in proteins and its use in analyzing interface evolution. Journal of Molecular Biology, 403(4):660–670, 2010.

[55] Rocco Moretti, Sarel J Fleishman, Rudi Agius, Mieczyslaw Torchala, Paul A Bates, Panagiotis L Kastritis, Joao PGLM Rodrigues, Mikaël Trellet, Alexandre MJJ Bonvin, Meng Cui, et al. Community-wide evaluation of methods for predicting the effect of mutations on protein–protein interactions. Proteins: Structure, Function, and Bioinformatics, 81(11):1980–1987, 2013.

[56] Sanghee Yoo, David G Myszka, Chin-yah Yeh, Maureen McMurray, Christopher P Hill, and Wesley I Sundquist. Molecular recognition in the HIV-1 capsid/cyclophilin A complex. Journal of Molecular Biology, 269(5):780–795, 1997.

[57] Theresa R Gamble, Felix F Vajdos, Sanghee Yoo, David K Worthylake, Megan Houseweart, Wesley I Sundquist, and Christopher P Hill. Crystal structure of human cyclophilin A bound to the amino-terminal domain of HIV-1 capsid. Cell, 87(7):1285–1294, 1996.

[58] Weizhong Li and Adam Godzik. Cd-hit: a fast program for clustering and comparing large sets of protein or nucleotide sequences. Bioinformatics, 22(13):1658–1659, 2006.

