## Supplementary Material for "Quantification of biases in predictions of protein-protein binding affinity changes upon mutations"

December 18, 2023

**4 Methods: Definition of additional mutations' subsets** **8**

**5 Dataset of mutually reverse mutations pairs** **8**

### 1 Error corrections from SKEMPI 1.0 to SKEMPI 2.0

The presence of errors in literature-based datasets (including SKEMPI 2.0) cannot be prevented. This leads to a certain degree of uncertainty in the collected values even if we suppose that the experimental conditions are always similar and that there are no fluctuations in the obtained experimental values.

This can be illustrated with the error corrections between SKEMPI 1.0 [1] and SKEMPI 2.0 [2] shown in Fig. S-1. Indeed, when we compare the entries that appear in the two sets, we observe some differences between their $\Delta\Delta G_b$ values. For example, the 16 mutations from [3] (occurring in human cyclophilin A bound to the amino-terminal domain of HIV-1 capsid (PDB 1AK4 [4])) have their $\Delta\Delta G_b$ values shifted by a factor 1.36 kcal/mol between SKEMPI 1.0 and SKEMPI 2.0. The difference is caused by an error of a factor 10 in the $K_D^{wt}$ values reported in SKEMPI 1.0.

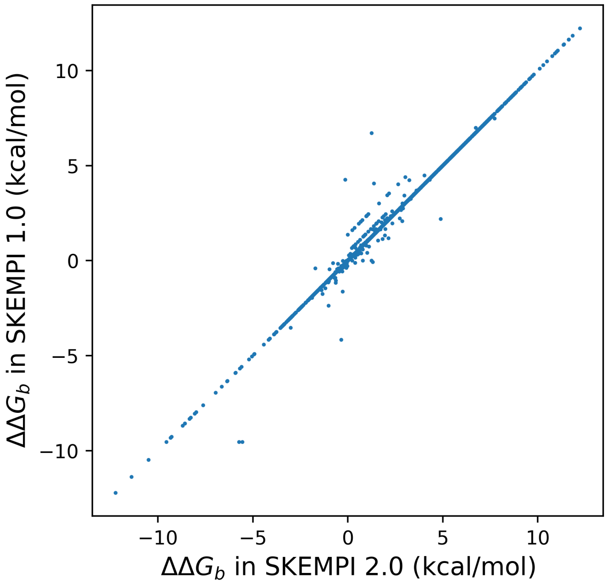

Figure S-1: Comparison of $\Delta\Delta G_b$ values (in kcal/mol) reported by SKEMPI 1.0 and SKEMPI 2.0.

### 2 Properties and biases in studied datasets

We have shown that our first benchmark dataset S2536 (based on SKEMPI 2.0 [2]) suffers from heavy unbalances in terms of mutation types, mutation location and positive shift of $\Delta\Delta G_b$ values. We note here that our structure quality filter does not impact these unbalances since S4193, the set of all non-redundant single-site mutations from SKEMPI 2.0, shows the same biases. Indeed, more that half of the mutations from S4193 are towards alanine, 181 mutation types occur less than 5 times and 67 mutation types are unrepresented. As for the mutations' location, 71% of the entries are mutations of the 8% residues located at the interface. Finally, the $\Delta\Delta G_b$ distribution from S4193 also shows a clear shift towards positive values (as shown in Table S-1 and Fig. S-2).

For the datasets deriving from the interaction of the RBD domain of the SARS-CoV-2 spike protein with ACE2 [5] (PDB structure 6M0J), we have also two datasets: C3684 containing all single-site mutations of all residues in 6M0J, and C380, our second benchmark dataset, containing only all single-site mutations at the protein-protein interface. Whether selecting only interface mutations or not, the $\Delta\Delta G_b$ distributions remain similar (Table S-1 and Fig. S-2), with the exception that C380 contains less neutral mutations (mutations with a $\Delta\Delta G_b$ value close to zero), which is expected for most mutations located far from the interface. The dataset C3684 as well as its subset C380 contain no stabilizing mutations and show an unusual peak around the maximal $\Delta\Delta G_b$ value (see Fig. S-2). This observation points to a minimal and a maximal experimental bound for the measured affinity values.

All datasets and corresponding $\Delta\Delta G_b$ experimental values are available at `https://github.com/3BioCompBio/DDGb_bias`.

Table S-1: Properties of the datasets S4193, S2536, C3684 and C380 and of their respective $\Delta\Delta G_b$ distributions (in kcal/mol).

| Dataset | #mutations | #complexes | Mean | Standard deviation | Minimum | Maximum |
|---|---|---|---|---|---|---|
| S4193 | 4193 | 319 | 0.97 | 1.74 | −12.22 | 12.22 |
| S2536 | 2536 | 205 | 1.11 | 1.99 | −12.22 | 12.22 |
| C3684 | 3684 | 1 | 1.03 | 1.42 | −0.30 | 4.84 |
| C380 | 380 | 1 | 1.24 | 1.25 | −0.30 | 4.80 |

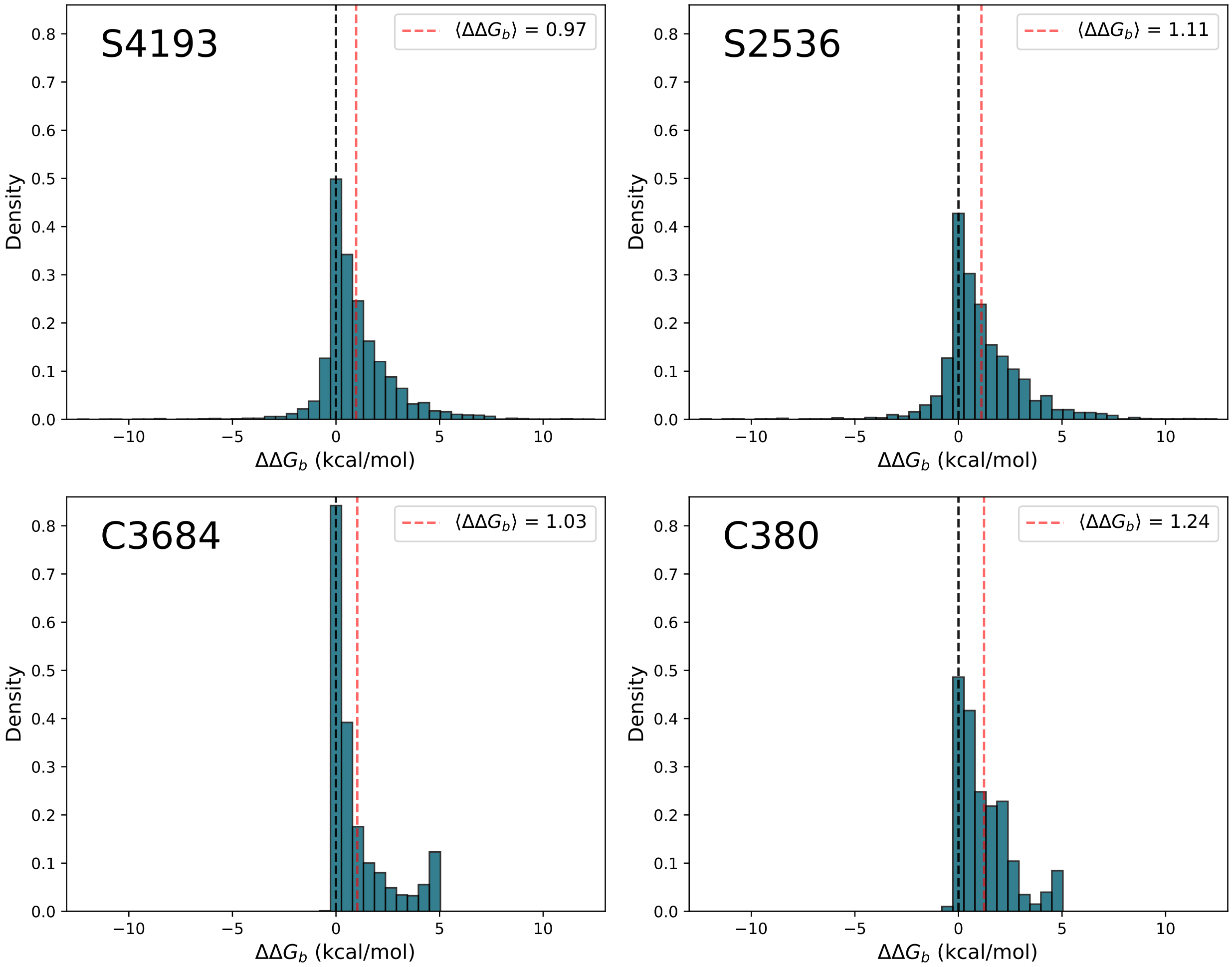

Figure S-2: Distributions of $\Delta\Delta G_b$ values (in kcal/mol) for S4193, S2536, C3684 and C380.

### 3 Binding affinity change predictions

In this section, we present all $\Delta\Delta G_b$ predictions of the eight benchmarked predictors (mCSM-PPI2 [6], MutaBind2 [7], BeAtMuSiC [8], SSIPe [9], SAAMBE-3D [10], NetTree [11], BindProfX [12] and FoldX [13]) on our two benchmark datasets S2536 (all single-site mutations on good-resolution crystal structures from SKEMPI 2.0) and C380 (all possible single-site mutations at the RBD-ACE2 interface [5]).

All datasets and predicted $\Delta\Delta G_b$ values are available at `https://github.com/3BioCompBio/DDGb_bias`.

#### 3.1 Comparison of predicted and experimental values

The scatter plots comparing the predicted and experimental $\Delta\Delta G_b$ values for the eight predictors on S2536 and C380 are shown in Figs S-3 and S-4.

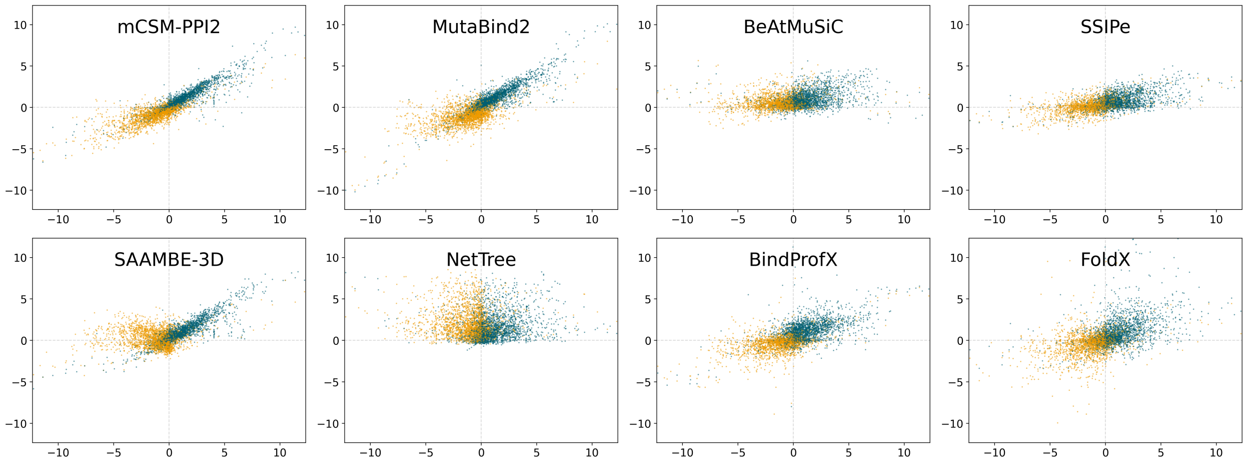

Figure S-3: Predicted $\Delta\Delta G_b$ values (in kcal/mol) as a function of experimental values on the dataset S2536. Direct mutations (S2536-D) are in blue and reverse mutations (S2536-R), in orange. Predictions are computed by mCSM-PPI2, MutaBind2, BeAtMuSiC, SSIPe, SAAMBE-3D, NetTree, BindProfX and FoldX.

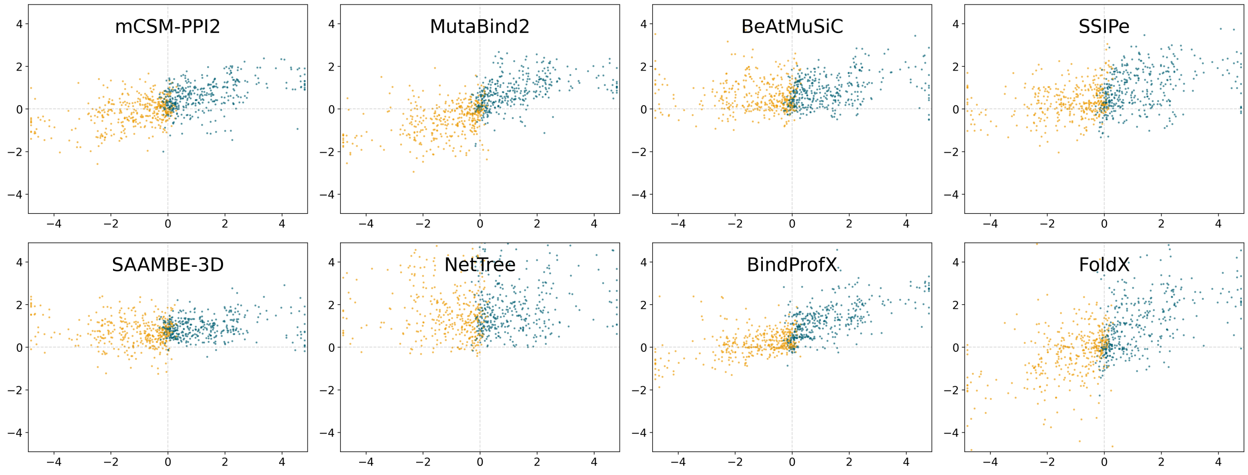

Figure S-4: Predicted $\Delta\Delta G_b$ values (in kcal/mol) as a function of experimental values on the dataset C380. Direct mutations (C380-D) are in blue and reverse mutations (C380-R), in orange. Predictions are computed by mCSM-PPI2, MutaBind2, BeAtMuSiC, SSIPe, SAAMBE-3D, NetTree, BindProfX and FoldX.

#### 3.2 Shift between direct and reverse mutations

The distributions of the shift $\delta = \Delta\Delta G_b^{wt\to mt} + \Delta\Delta G_b^{mt\to wt}$ between predictions on direct and reverse mutations are represented for the eight benchmarked predictors in Fig. S-5 for the dataset S2536 and in Fig. S-6 for C380. Note that theoretically, the shift is supposed to be zero. Even if some fluctuations are unavoidable for a predictor, a deviation of the mean shift is a measure of its bias towards stabilizing mutations. We consider the shifts on C380 as more reliable than on S2536 since its mutations are systematic and unknown to the eight predictors.

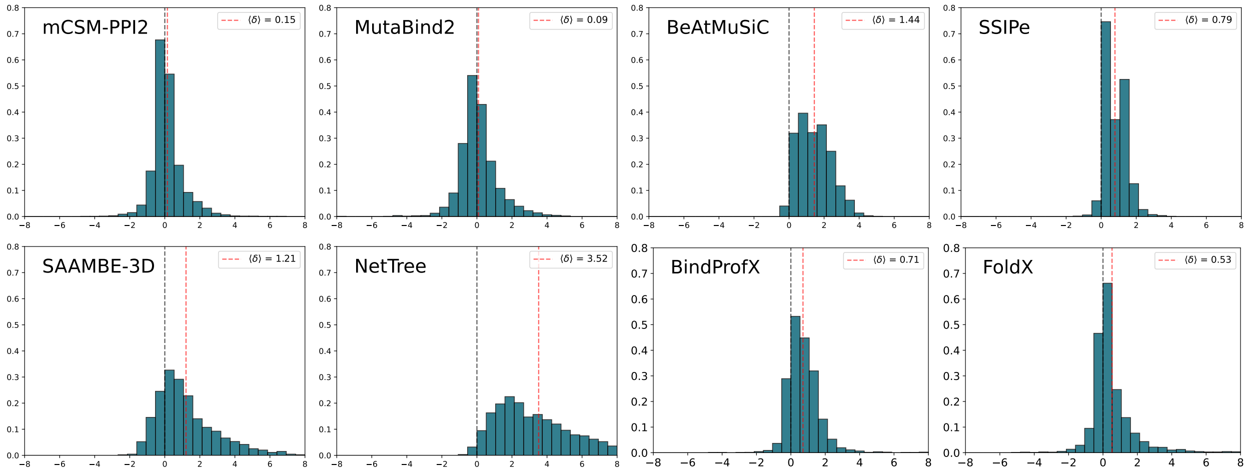

Figure S-5: Distributions of the $\Delta\Delta G_b$ shift $\delta$ (in kcal/mol) between direct mutations from S2536-D and reverse mutations from S2536-R for mCSM-PPI2, MutaBind2, BeAtMuSiC, SSIPe, SAAMBE-3D, NetTree, BindProfX and FoldX.

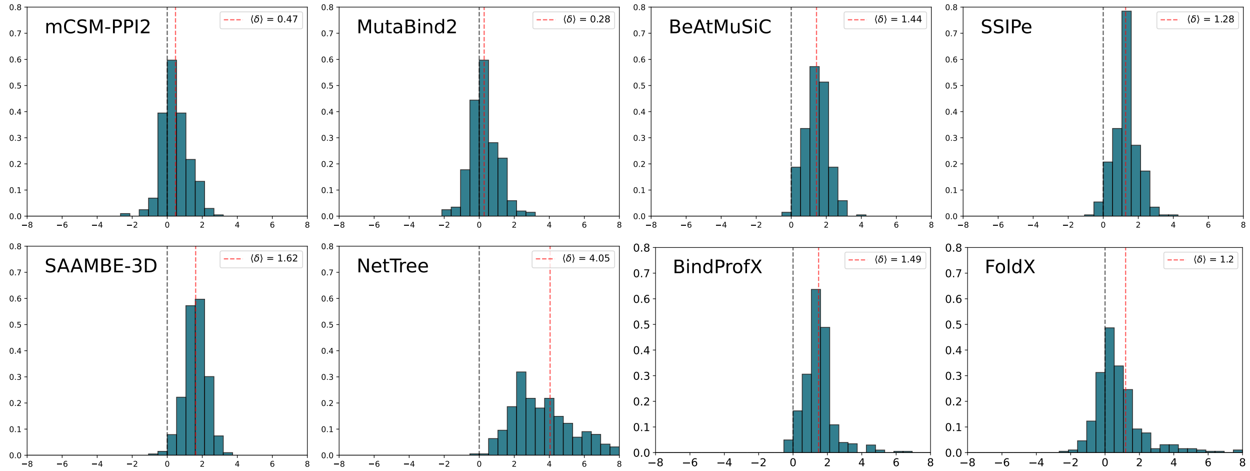

Figure S-6: Distributions of the $\Delta\Delta G_b$ shift $\delta$ (in kcal/mol) between direct mutations from C380-D and reverse mutations from C380-R for mCSM-PPI2, MutaBind2, BeAtMuSiC, SSIPe, SAAMBE-3D, NetTree, BindProfX and FoldX.

#### 3.3 Performances of the predictions

Fig. S-7 shows the performances of the eight benchmarked predictors on S2536 and C380 for direct and reverse mutations measured by the Pearson correlation ($\rho$), the Spearman rank correlation (r) and the root mean squared error (RMSE). In addition, this figure shows the performances of the symmetrized versions of the eight predictors, which are computed from the predictions on both the direct mutation and corresponding reverse mutation:

$$\overline{\Delta\Delta G_b}^{wt\to mt} := \frac{\Delta\Delta G_b^{wt\to mt} - \Delta\Delta G_b^{mt\to wt}}{2},$$
$$\overline{\Delta\Delta G_b}^{mt\to wt} := \frac{\Delta\Delta G_b^{mt\to wt} - \Delta\Delta G_b^{wt\to mt}}{2}.$$

Note that by its definition, the symmetrized version of a predictor has the exact same performances on direct and on reverse mutations. Thus, the performances of the symmetrized predictors are indistinguishably represented for both direct and reverse datasets.

All numeric values of these metrics (as well as of other metrics, i.e. the Kendall rank correlation, normalized root mean squared error, mean squared error, mean absolute error and mean signed error) are available at `https://github.com/3BioCompBio/DDGb_bias`.

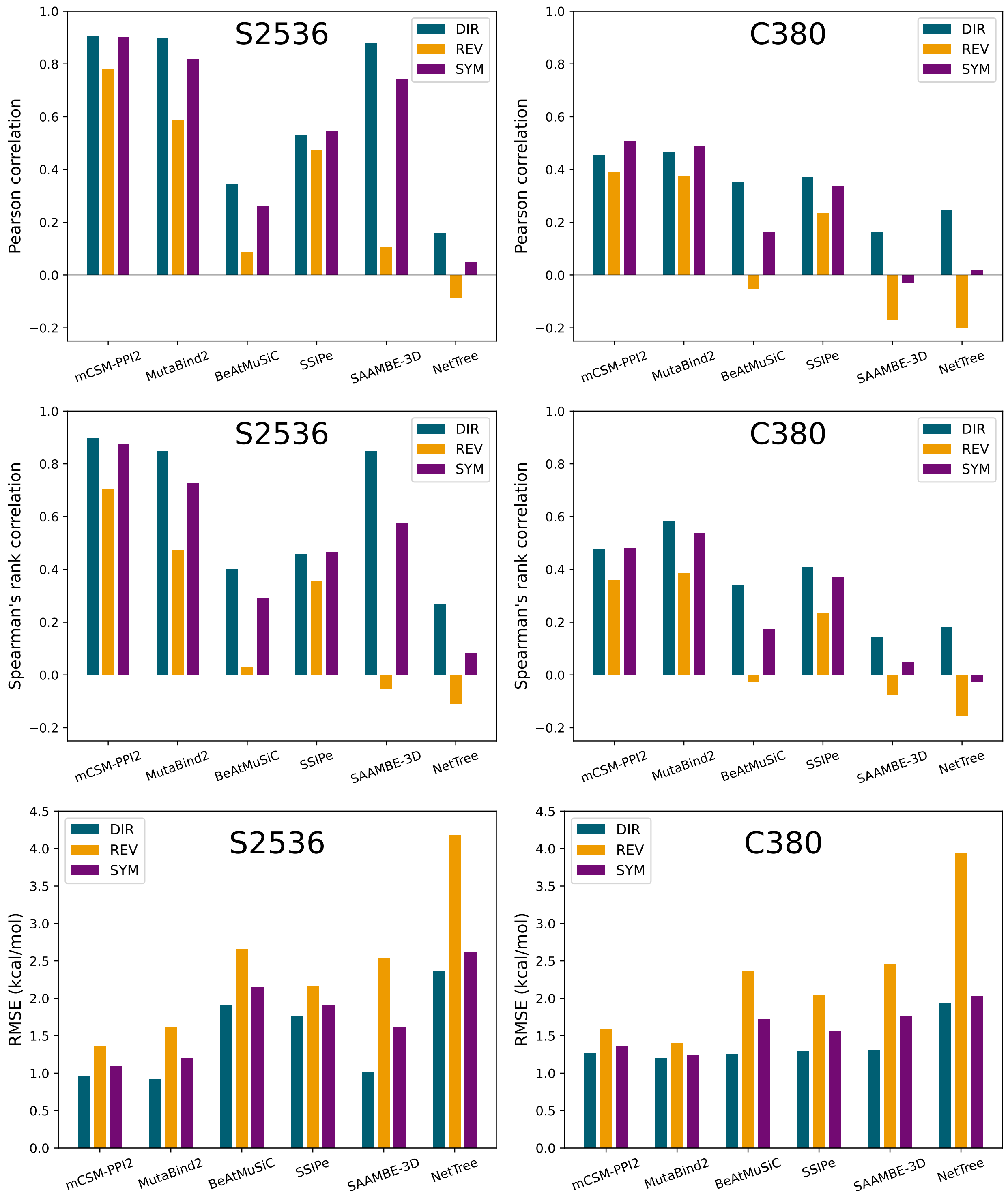

Figure S-7: Performances of the eight benchmarked predictors (mCSM-PPI2, MutaBind2, BeAtMuSiC, SSIPe, SAAMBE-3D, NetTree, BindProfX and FoldX) on the datasets S2536 and C380 for direct mutations (in blue), for reverse mutations (in orange) and for the symmetrized predictors (in purple). The performances are measured using the Pearson correlation, the Spearman rank correlation and the RMSE (in kcal/mol).

### 4 Methods: Definition of additional mutations' subsets

As for the RSA, we used the MuSiC suite [14, 15] to assign the secondary structure of a residue in a 3D structure. We considered three categories: helix (A) grouping $\alpha$- and $3_{10}$-helices, sheet (B) grouping isolated and extended $\beta$s, and other (X).

For the solvent exposure, we used the cRSA (RSA of the residue in the complex) to distinguish two categories: buried residues (BUR; cRSA $< 30\%$) and exposed residues (EXP; cRSA $\geq 30\%$).

Finally, we also used cRSA, iRSA (RSA computed in the structure containing only one interactant) and $\Delta$RSA := iRSA − cRSA values to classify each residue into one of the five structural regions defined in [16]: interior (INT; $\Delta$RSA $\leq 5\%$ and cRSA $\leq 25\%$ ), surface (SUR; $\Delta$RSA $\leq 5\%$ and cRSA $> 25\%$ ), support (SUP; $\Delta$RSA $> 5\%$ and iRSA $\leq 25\%$), core (COR; $\Delta$RSA $> 5\%$, iRSA $> 25\%$ and cRSA $\leq 25\%$) and rim (RIM; $\Delta$RSA $> 5\%$ and cRSA $> 25\%$). Regions SUP, RIM and COR subdivide the interface while INT and SUP are the two non-interface regions.

Performances of the eight benchmarked predictors on all subsets of S2536-D are available at `https://github.com/3BioCompBio/DDGb_bias`).

### 5 Dataset of mutually reverse mutations pairs

We constructed the symmetric dataset SK2$^{sym}$ containing the 207 pairs of complexes from SKEMPI 2.0 which are separated by a single mutation. We used for that purpose the software CD-HIT [17] to fasten the all-to-all comparison of chains' sequences. The SK2$^{sym}$ dataset has the advantage that both the "wild-type" and "mutant" complexes have an experimentally resolved PDB structure and thus $\Delta\Delta G_b$ predictions of the "direct" and the "reverse" mutations do not require a modeled structure. Note that some pairs of complexes have some minor additional differences (other than the single-site mutation that separates them). These differences are specified in the dataset. The dataset is available at `https://github.com/3BioCompBio/DDGb_bias`).

### References

[1] Iain H Moal and Juan Fernández-Recio. SKEMPI: a structural kinetic and energetic database of mutant protein interactions and its use in empirical models. *Bioinformatics*, 28(20):2600–2607, 2012.

[2] Justina Jankauskaitė, Brian Jiménez-García, Justas Dapkūnas, Juan Fernández-Recio, and Iain H Moal. SKEMPI 2.0: an updated benchmark of changes in protein–protein binding energy, kinetics and thermodynamics upon mutation. *Bioinformatics*, 35(3):462–469, 2019.

[3] Sanghee Yoo, David G Myszka, Chin-yah Yeh, Maureen McMurray, Christopher P Hill, and Wesley I Sundquist. Molecular recognition in the HIV-1 capsid/cyclophilin A complex. *Journal of Molecular Biology*, 269(5):780–795, 1997.

[4] Theresa R Gamble, Felix F Vajdos, Sanghee Yoo, David K Worthylake, Megan Houseweart, Wesley I Sundquist, and Christopher P Hill. Crystal structure of human cyclophilin A bound to the amino-terminal domain of HIV-1 capsid. *Cell*, 87(7):1285–1294, 1996.

[5] Tyler N Starr, Allison J Greaney, Sarah K Hilton, Daniel Ellis, Katharine HD Crawford, Adam S Dingens, Mary Jane Navarro, John E Bowen, M Alejandra Tortorici, Alexandra C Walls, et al. Deep mutational scanning of SARS-CoV-2 receptor binding domain reveals constraints on folding and ACE2 binding. *Cell*, 182(5):1295–1310, 2020.

[6] Carlos HM Rodrigues, Yoochan Myung, Douglas EV Pires, and David B Ascher. mCSM-PPI2: predicting the effects of mutations on protein–protein interactions. *Nucleic Acids Research*, 47(W1):W338–W344, 2019.

[7] Ning Zhang, Yuting Chen, Haoyu Lu, Feiyang Zhao, Roberto Vera Alvarez, Alexander Goncearenco, Anna R Panchenko, and Minghui Li. MutaBind2: predicting the impacts of single and multiple mutations on protein-protein interactions. *Iscience*, 23(3):100939, 2020.

[8] Yves Dehouck, Jean Marc Kwasigroch, Marianne Rooman, and Dimitri Gilis. BeAtMuSiC: prediction of changes in protein–protein binding affinity on mutations. *Nucleic Acids Research*, 41(W1):W333–W339, 2013.

[9] Xiaoqiang Huang, Wei Zheng, Robin Pearce, and Yang Zhang. SSIPe: Accurately estimating protein–protein binding affinity change upon mutations using evolutionary profiles in combination with an optimized physical energy function. *Bioinformatics*, 36(8):2429–2437, 2020.

[10] Swagata Pahari, Gen Li, Adithya Krishna Murthy, Siqi Liang, Robert Fragoza, Haiyuan Yu, and Emil Alexov. SAAMBE-3D: predicting effect of mutations on protein–protein interactions. *International Journal of Molecular Sciences*, 21(7):2563, 2020.

[11] Menglun Wang, Zixuan Cang, and Guo-Wei Wei. A topology-based network tree for the prediction of protein–protein binding affinity changes following mutation. *Nature Machine Intelligence*, 2(2):116–123, 2020.

[12] Peng Xiong, Chengxin Zhang, Wei Zheng, and Yang Zhang. BindProfX: assessing mutation-induced binding affinity change by protein interface profiles with pseudo-counts. *Journal of Molecular Biology*, 429(3):426–434, 2017.

[13] Joost Schymkowitz, Jesper Borg, Francois Stricher, Robby Nys, Frederic Rousseau, and Luis Serrano. The FoldX web server: an online force field. *Nucleic Acids Research*, 33(suppl_2):W382–W388, 2005.

[14] Georgios A Dalkas, Fabian Teheux, Jean Marc Kwasigroch, and Marianne Rooman. Cation–$\pi$, amino–$\pi$, $\pi$–$\pi$, and H-bond interactions stabilize antigen–antibody interfaces. *Proteins: Structure, Function, and Bioinformatics*, 82(9):1734–1746, 2014.

[15] Yves Dehouck, Aline Grosfils, Benjamin Folch, Dimitri Gilis, Philippe Bogaerts, and Marianne Rooman. Fast and accurate predictions of protein stability changes upon mutations using statistical potentials and neural networks: PoPMuSiC-2.0. *Bioinformatics*, 25(19):2537–2543, 2009.

[16] Emmanuel D Levy. A simple definition of structural regions in proteins and its use in analyzing interface evolution. *Journal of Molecular Biology*, 403(4):660–670, 2010.

[17] Weizhong Li and Adam Godzik. Cd-hit: a fast program for clustering and comparing large sets of protein or nucleotide sequences. *Bioinformatics*, 22(13):1658–1659, 2006.
